# Venom protection by broadly neutralizing antibody from a snakebite subject

**DOI:** 10.1101/2022.09.26.507364

**Authors:** Jacob Glanville, Joel Christian Andrade, Mark Bellin, Sangil Kim, Sergei Pletnev, David Tsao, Raffaello Verardi, Rishi Bedi, Sindy Liao, Raymond Newland, Nicholas L. Bayless, Sawsan Youssef, Ena Tully, Baoshan Zhang, Tatsiana Bylund, Sujeong Kim, Tracy Liu, Peter D. Kwong

## Abstract

Snake envenomation is a neglected tropical disease, causing >100,000 deaths and 300,000 permanent disabilities in humans annually. Could monoclonal antibody technology provide a solution? Here, we recover Centi-3FTX-D09, a potent broadly neutralizing antivenom antibody from the B-cell memory of a human subject with snake venom exposure. Centi-3FTX-D09 recognized a conserved neutralizing epitope on long 3-finger toxins (3FTXs), a dominant snake neurotoxin. Crystal structures of Centi-3FTX-D09 in complex with 3FTXs from mamba, taipan, krait, and cobra revealed epitope mimicry of the interface between these neurotoxins and their host target, the nicotinic acetylcholine receptor. Centi-3FTX-D09 provided *in-vivo* protection against diverse recombinant long 3FTXs, *in-vivo* rescue from whole venom challenge from cobras, black mamba, and king cobra, and, when combined with the phospholipase inhibitor varespladib, *in-vivo* protection extending to a majority of tested elapid venoms. Thus, a single antibody can broadly neutralize long neurotoxins and contribute to broad protection from envenomation.

## Introduction

In 2018, snakebite envenoming became ranked in the World Health Organization’s list of neglected tropical diseases^1^, every year causing 81,000-138,000 deaths and leaving another 300,000-400,000 permanently disabled^2,3^. For over a century, the standard of care for snakebite envenoming has been antivenom: a polyclonal serum preparation derived from animals immunized with venom from one or more snakes^4^. While effective, antivenom serotherapies present several challenges. Treatment with non-human antibodies often results in early-onset adverse reactions in patients, which can lead to more severe reactions if the antivenom were to be ever used again in the same patient^5^. Envenomed patients typically require five or more doses of antivenom, as polyclonal sera-derived Fab formulations have short half-lives, are contaminated with 5-22% non-antibody proteins, and have been reported to contain only 9-15% of venom-specific antibodies to any given snake in the case of polyvalent polyclonal antivenoms^6–8^. Moreover, antivenom developed for a single species requires the correct identification of the specific snake that bit the victim, which is often not possible for victims or healthcare workers not well-versed in snake phenotypes^6,9^. Although a global public health problem, research into improved envenoming treatments is hampered by the heterogeneity of venom toxins across species, the relatively low market value of a disease primarily afflicting the developing world, and the great fracturing of that market by differing requirements of the many different species-restricted antivenoms^10^.

Historical efforts to isolate therapeutic antibodies targeting individual toxins have also shown limited and species-specific efficacy. Hybridomas made from B cells of mice immunized with short-chain neurotoxin allowed for isolation of a broadly-reactive anti-neurotoxin monoclonal antibody, but *in-vivo* efficacy was never evaluated^11^. Antibody engineering approaches designed to artificially mimic the nicotinic receptor binding site have also resulted in antibody clones targeting highly species-specific epitopes^12^. More recently, high-throughput methods have been employed to discover anti-venom antibodies, including an anti-cobra antibody discovered using a phage display library of naïve single-chain variable fragments (scFv)^13^. Recently, an anti-3FTX antibody with detectable binding extending to multiple genera have been reported, but with full in-vivo protection limited to a single cobra species, with only partial rescue protection against the single cobra species, when the antibody was administered 10 minutes following venom injection^14^.

Although desirable, a single broad-spectrum, fully-human, antivenom has never been developed, predominantly due to complexity of snake venom and diversity of snake species. There are over 500 genetically diverse venomous snake species globally that span 167 million years of Toxicofera evolution^15^. Each snake produces 5-70 toxins, or unique proteins, in their venom, and there is abundant polymorphism of venom proteins even between individuals of a single snake species^16^. Thus, even if an antibody were discovered for every toxin of every venomous snake, it would not be possible to combine all of these antibodies into a single antivenom, as each antibody would be at far too low of a dose to be therapeutically relevant^17–19^.

In this study, we sought to determine whether broadly neutralizing antibodies could protect mice against entire classes of snake toxins as well as whole venom challenge. We chose to begin with α-neurotoxin long 3-finger toxins (3FTXs), the dominant neurotoxin of the elapid family of snakes^20–22^. The elapid family of snakes account for ∼60% of venomous snake species and includes over 50 genera, 300 species, and over 170 subspecies^23^. The long three-finger toxins (3FTXs) are amongst the deadliest toxins in elapid venom^24^. Long 3FTX orthologs are found in nearly all elapid neurotoxic snakes, and through highly diverse (**Supplementary Fig. 1**), they bind with high affinity to both muscle and neuronal nicotinic acetylcholine receptors (nAChR) and impair neuromuscular and neuronal transmission^25^. The extreme conservation of the nAChR/acetylcholine interface across all vertebrates enables a single toxin to productively paralyze neurons from mammals, birds, reptiles, amphibians, and fish (**Supplementary Fig. 2**). This evolutionary constraint also requires that long 3FTXs from all snake species maintain a conserved interface with the nAChR^26^, thereby presenting a plausible target for broadly neutralizing anti-3FTX antibodies. Here we report the isolation of such a broadly neutralizing antibody, and characterization of *in-vivo* protection by this antibody alone and in combination with varespladib^27^, a small-molecule PLA2 inhibitor.

### *In-vivo* and *in-vitro* selection for broadly neutralizing 3FTx antibodies

Broadly neutralizing antibodies (bnAbs) against snake neurotoxins were isolated from an adult male subject with an extensive history of diverse snake venom exposure (**Fig. 1a)**.

**Figure 1.**
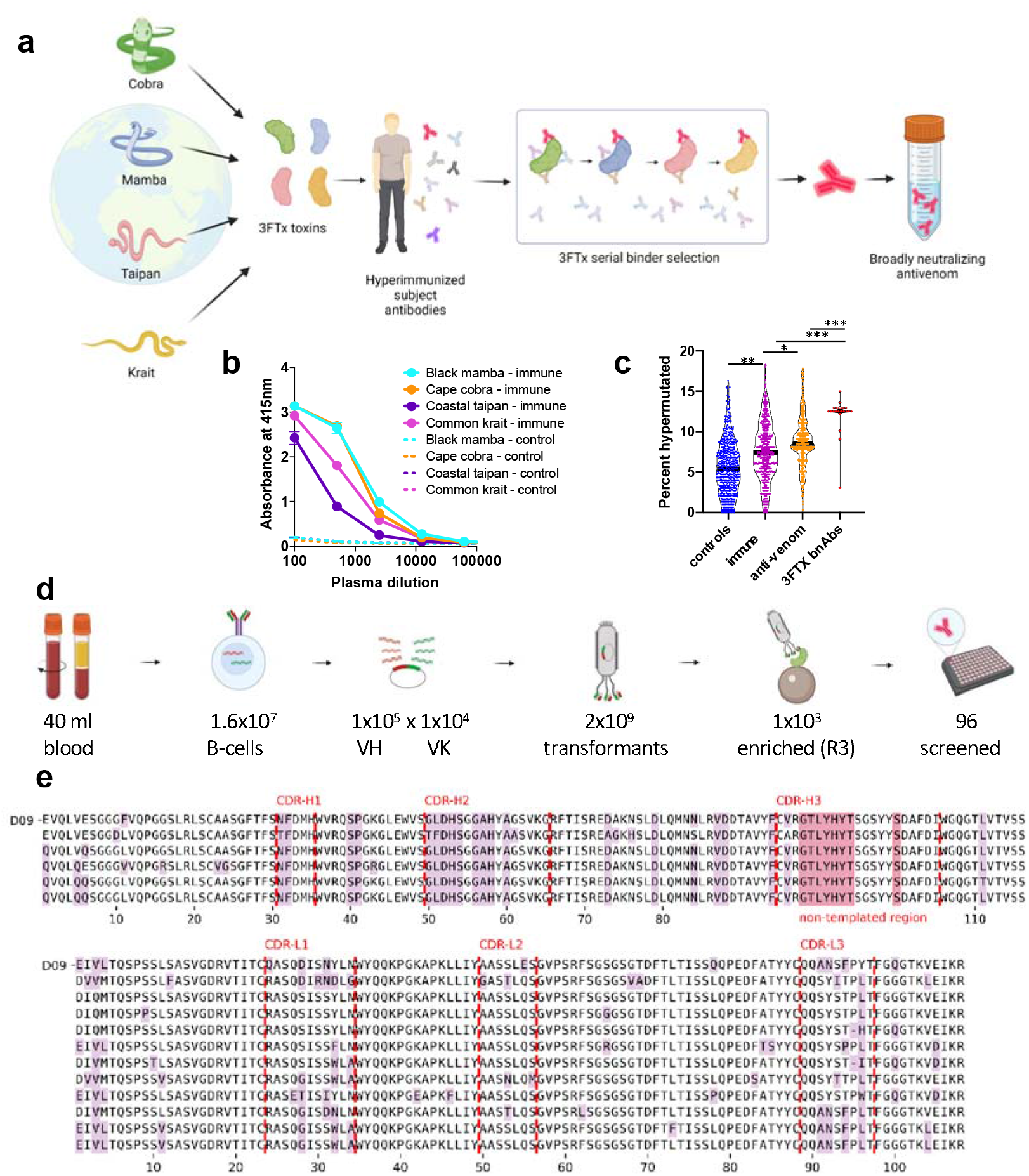
In-vivo and in-vitro selection for broadly neutralizing long 3FTX antibodies. **(a)** Immune library from hyperimmune subject was sequentially panned against recombinant long 3FTX neurotoxins from cobra, taipan, mamba and krait. **(b)** Serum reactivity to recombinant long 3FTX neurotoxins, hyperimmune subject versus venom-naive donors. **(c)** Antibody repertoire somatic hypermutation. **(d)** Down-selection and screening of 16 million B cells to identify final lead D09 antibody lineage. **(e)** D09 lineage mutational variants for confirmed binders of **(top**) heavy and (**bottom)** light chain Fv domains (mutations from germline in purple; VDJ non-templated base positions in salmon.

A hyper-immune human subject was identified with a documented 18-year history of 756 immunizations to venoms spanning 2001 to 2018. Documented immunizations included mambas (*D. polylepis, D. viridis, D. angusticeps, D. jamesoni*), cobras (*N. kaouthia, N. haje, N. melanoleuca, N. nivea*), rattlesnakes (*C. atrox, C. scutulatus*), water cobras (*N. annulata, N. cristyi*), taipans (*O. scutellatu*s, *O. scutellatus canni*), as well as *M. fulvius* (coral snake), *B. caeruleus* (common krait), *B. multicinctus* (banded krait), *N. scutatus* (tiger snake), and *P. textilis* (eastern brown snake). Hypothesizing that this repeated cyclical pattern of diverse venom exposure may have selected for broadly-reactive antivenom antibodies that recognized conserved epitopes shared across the venom toxins of multiple snake species, we sought to isolate such bnAbs from this subject (see **Ethics Statement**).

In a non-interventional study design, two 20 ml samples of blood were collected during the subject’s standard self-immunization schedule: one 20 ml collection following a three-month venom immunization holiday (Day 0), and the second 20 ml collection 7 days following the end of the subject’s 21-day routine self-immunization schedule with black mamba, western diamondback, and coastal taipan (Day 28). Informed consent was obtained for blood sample collection, and collection was conducted in accordance with Western Institutional Review Board IRB Exemption Work Order #1-1209200-1 (**Ethics Statement**). Blood samples were separated into plasma and PBMCs (see **Online Methods:Plasma & PBMCs**). Relative to snake venom-naive healthy control plasma, significantly elevated antibodies were detected in hyperimmune subject plasma against a panel of long 3FTX from black mamba, cape cobra, coastal taipan, and common krait to a level of 1:10,000 dilution (**Fig. 1b, Online Methods:Serum ELISA**). From the PBMCs, B-cell receptor repertoire DNA was isolated from each blood sample, and the isolated repertoires underwent high-throughput sequencing (**Supplementary Table 1; Online Methods:Amplicons**). The total VH repertoire somatic hypermutation (SHM) rate was elevated in the hyperimmune subject at Day 0 (7.32 +/- 2.69) compared with a panel of healthy adult controls (5.66 +/- 2.52) (**Fig. 1c; Online Methods:Informatics**). The SHM rate increased at Day 28 following the immunization schedule (7.65 +/- 2.61). The SHM rate was further significantly elevated in the venom-specific repertoire of the subject (8.61 +/- 2.23), obtained by enrichment panning against biotinylated whole venom from 4 snake species. The SHM rate was highest in the 64 broadly cross-reactive antivenom antibodies isolated against homologs of long 3FTX (12.06 +/- 1.62). Thus, the hyperimmune subject appeared to have an elevated total SHM profile compared with healthy controls, this elevated SHM profile was enriched in the antivenom-specific repertoire, and particularly elevated in isolated broadly neutralizing antibodies.

From the hyperimmune subject’s PBMCs, broadly neutralizing anti-long 3FTX monoclonal antibodies were isolated by phage display **(Fig. 1, d and e)**. From 40 ml of blood, containing approximately 16 million B cells, antibody Fv VH and VL domains were amplified by multiplex phage adaptor PCR (**Supplementary Table 1; Online Methods:Amplicons**). Sequencing of 4.3 million amplicons identified at least 45,174 unique CDR-H3 VH sequences and 36,817 unique CDR-L3 sequences, excluding singletons. These domains were assembled combinatorially into an m13 VH-VL scFv-pIII fusion display vector and transformed to a final library size of 2.21e9, such that every observed VH domain was associated with every VL domain in the library, including every original native VH/VL pair (**Online Methods:Library**).

To isolate broadly reactive antibodies from the hyperimmune subject antibody library, the library was sequentially panned against four recombinant α-neurotoxin ortholog representatives from four diverse genera of elapidae. The four orthologs chosen were alpha-elapitotoxin-Dpp2a (3L24_DENPO) from *D. polylepis* (black mamba) of Sub-Sarahan Africa, alpha-cobratoxin/long neurotoxin 1 (3L21_NAJNI) from *N. nivea* (Cape cobra) of Southern Africa, Long neurotoxin 1 (3L21_OXYSC) from *O. scutellatus scutellatus* (coastal taipan) from Australia, and Alpha-delta bungarotoxin (3L2A_BUNCE) from *B. caeruleus* (common krait) from Asia. Evolutionarily, these four distinct genera of elapids span 40 million years since most recent common ancestor, with highly diverse long 3FTX orthologs that share only 48-64% identity at the amino acid level (**Fig. S1**). Clinically, they represent some of the deadliest elapids to humans. The four recombinant α-neurotoxin orthologs were expressed with C-terminal site-specific biotinylation avitags in HEK293 cell culture, tag-purified, and lethal neurotoxic functional activity was confirmed *in-vivo* in C57BL/6 mouse Maximum Tolerated Dose (MTD) studies (**Online Methods:Recombinants**).

Four sequential rounds of soluble-phase automated panning were performed, using a different ortholog of long 3FTX in each subsequent round to select for breadth (**Fig. 1a, Online Methods:Panning**). Following the third round, Illumina MiSeq repertoire sequencing was performed, observing approximately one thousand unique enriched VH clonal variants (**Fig. 1d**), as well as a significant enrichment of clones with more somatic hypermutation than observed in the hyperimmune subject’s total repertoire (**Fig. 1c**). Following the fourth round of selection, 376 clones were isolated and screened for reactivity to the cobra, taipan, krait, and mamba long 3FTX by ELISA (**Fig. 1d; Supplementary Table 2; Online Methods:PPE Screening**). Sixty-four broadly reactive neurotoxin-specific clones were identified, and upon sequencing 61 of the 64 (95%) were found to be from a single dominant lineage D09 that utilized the same VDJ rearrangement and contained the clone Centi-3FTX-D09 as a member (**Fig. 1e; Online Methods:Informatics**). The D09 lineage utilized IGHV3-13 V-gene segment, a long (19 amino acids) CDR-H3 loop with a 7 amino acid non-templated V-D junctional region, and bore evidence of extensive somatic hypermutation, with 22 amino acid mutations relative to germline (77.4% identity) in the heavy chain. The mutations were concentrated in the CDRs, with 9 mutations in CDR-H2, 2 mutations in CDR-H1, and 1 mutation and 8 non-templated amino acids in CDR-H3 (**Fig. 1e**). Six definitive heavy chain SHM variants were identified, all showing evidence of extensive SHM and a common origin from a single parental B-cell clone. The light chain had significant restriction to IGKV1-39 (85%), with SHM mutations including 3 in CDR-L1, 1 in CDR-L2, 4 putative non-templated base mediated amino acids in CDR-L3, and 4 mutations in the frameworks. Neither germline reversion of all 8 framework SHM mutations in the VH domain nor all 4 framework SHM mutations in the VK domain impacted binding, and both Centi-3FTX-D09 and germlined D09 lineage variants exhibited unusually high thermostability and aggregation resistance (75 °C Tm1, 75 °C Tagg 266).

### Affinity and breadth of Centi-3FTX-D09

Initial ELISA screening indicated broad reactivity of lineage D09 to the representative recombinant long 3FTX neurotoxins from cape cobra, common krait, coastal taipan, and black mamba (**Supplementary Table 2**). Kinetics assays using Surface Plasmon Resonance (SPR) performed on Biacore 8K demonstrated picomolar affinity monovalent interactions between lineage member Centi-3FTX-D09 and long 3FTX neurotoxins from diverse species: <74 pM to black mamba, <37 pM to Cape cobra, and 490 pM to coastal taipan, as well as high affinity interactions to these toxins by Centi-3FTX-B11, another member of the lineage (**Fig. 2a-c; Supplementary Fig. 3; Online Methods:Kinetics**).

**Figure 2.**
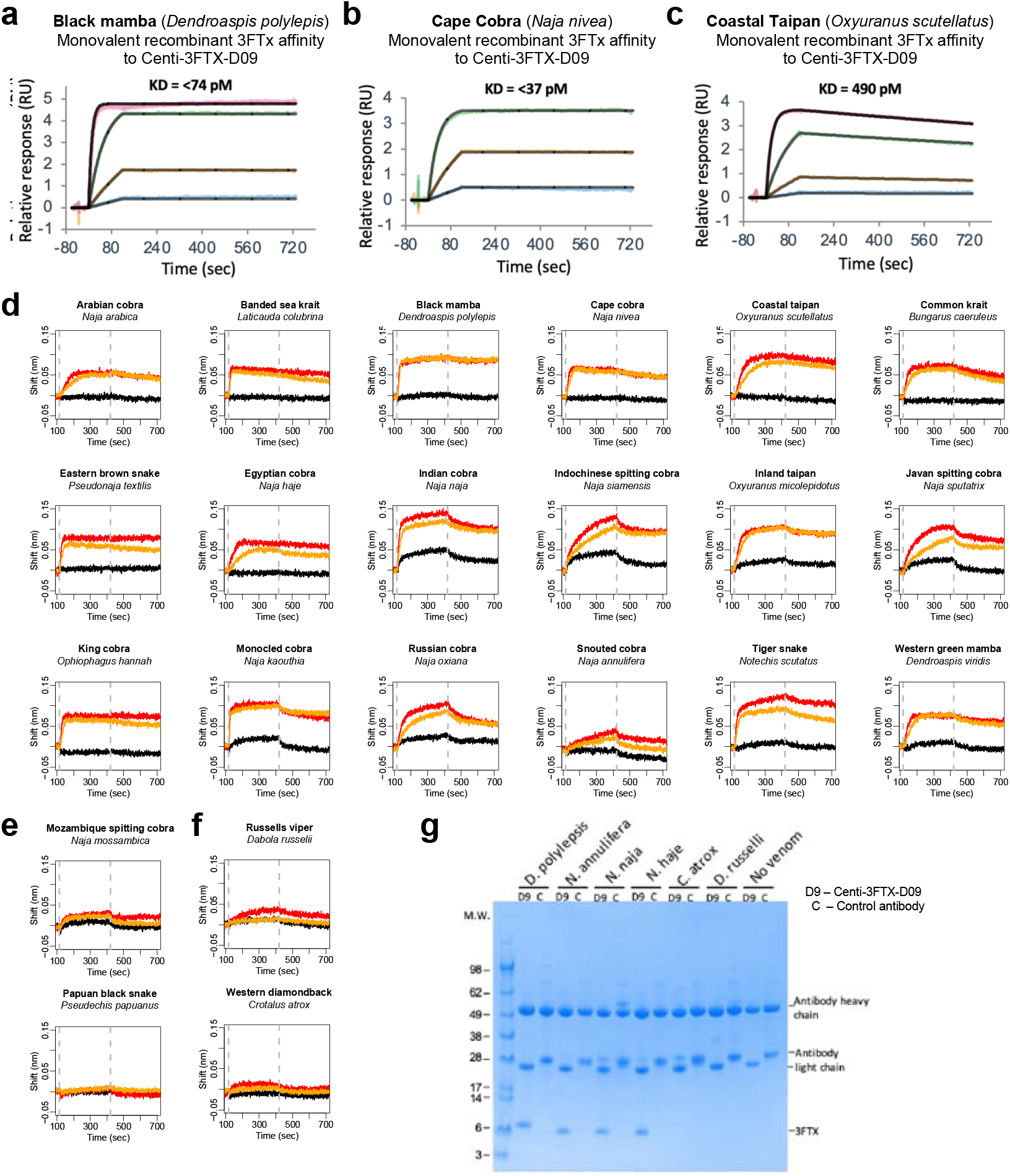
Affinity and breadth of Centi-3FTX-D09. **(a-c)** Monovalent affinity by Biacore 8K surface plasmon resonance kinetics of Centi-3FTX-D09 vs monovalent recombinant long 3FTX for **(a)** black mamba, **(b)** cape cobra, and **(c)** coastal taipan. **(d-f)** Whole venom Biolayer Iinterferometry reactivity of Centi-3FTX-D09 versus **(d)** eighteen elapid species with confirmed reactivity; **(e)** two elapid species with no/marginal reactivity, and **(f)** two negative control viperid venoms with no known 3FTX (black: off-target mAb vs venom at 500nM, Red: Centi-3FTX-D09 vs venom 500nM, orange: Centi-3FTX-D09 vs venom 250nM) **(g)** Immunoprecipitation by Centi-3FTX-D09 of long 3FTXs from whole venom (3FTX identity confirmed by mass-spectrometry as detailed in **Supplementary Table 3**).

To further characterize the breadth of Centi-3FTX-D09 reactivity, we used biolayer interferometry to assess the reactivity of Centi-3FTX-D09 to whole venom from 20 elapid species. In 18 species, we observed clear reactivity (**Fig. 2d**) and, in two, *N. mossambica* and *P. papuanus*, we observed no or marginal reactivity (**Fig. 2e**). Consistent with this observation, *P. papuanus* venom has been reported to contain no long 3FTX^28^, and for both species, no long neurotoxin 3FTX family members were found in Uniprot. As negative controls, we also tested two viperid venoms not expected to contain long 3FTX (from *D. russelii* and *C. atrox)* and observed no/marginal reactivity (**Fig. 2f**).

To confirm the reactivity in venom corresponding to recognition by Centi-3FTX-D09 of a long 3FTX, we performed immunoprecipitation pull-down and mass-spectrometry peptide sequencing experiments with Centi-3FTX-D09 and whole venom and observed clear evidence for long 3FTX binding with bands in the 6-7kDa range observed for *D. polylepsis, N. naja, N. annulifera*, and *N. haje*, but not with *D. russelii, C. atrox* (**Fig. 2g)**. Mass-spectrometry confirmed the specific molecular identity of the pulled-down peptides to be long 3FTX in every instance where a band was observed (**Supplementary Table 3**).

Overall, these binding experiments demonstrate high affinity of Centi-3FTX-D09, with breadth extending to most long 3FTXs.

### Crystal structures of Centi-3FTX-D09 with long 3FTXs reveal similarity in recognition between antibody and acetylcholine receptor

To provide an atomic-level explanation for the broad reactivity of Centi-3FTX-D09 with long 3FTXs, we determined crystal structures of the antigen-binding fragment (Fab) of Centi-3FTX-D09 in complex with long 3FTXs from cape cobra, common krait, coastal taipan, and black mamba (**Fig. 3a** and **Supplementary Table 4**). Analysis of the interfaces between Centi-3FTX-D09 Fab and long 3FTXs from these diverse snake genera revealed buried surfaces areas of ∼1600 Å^2^, shared between Fab and toxin (**Supplementary Tables 5 and 6**). The interactive surface area was contributed primarily by the heavy chain 3^rd^ complementarity determining region (CDR H3), although CDR H1 and L1 and L2 contributed a mixture of hydrophobic interactions and hydrogren bonds (H-bonds).

**Figure 3.**
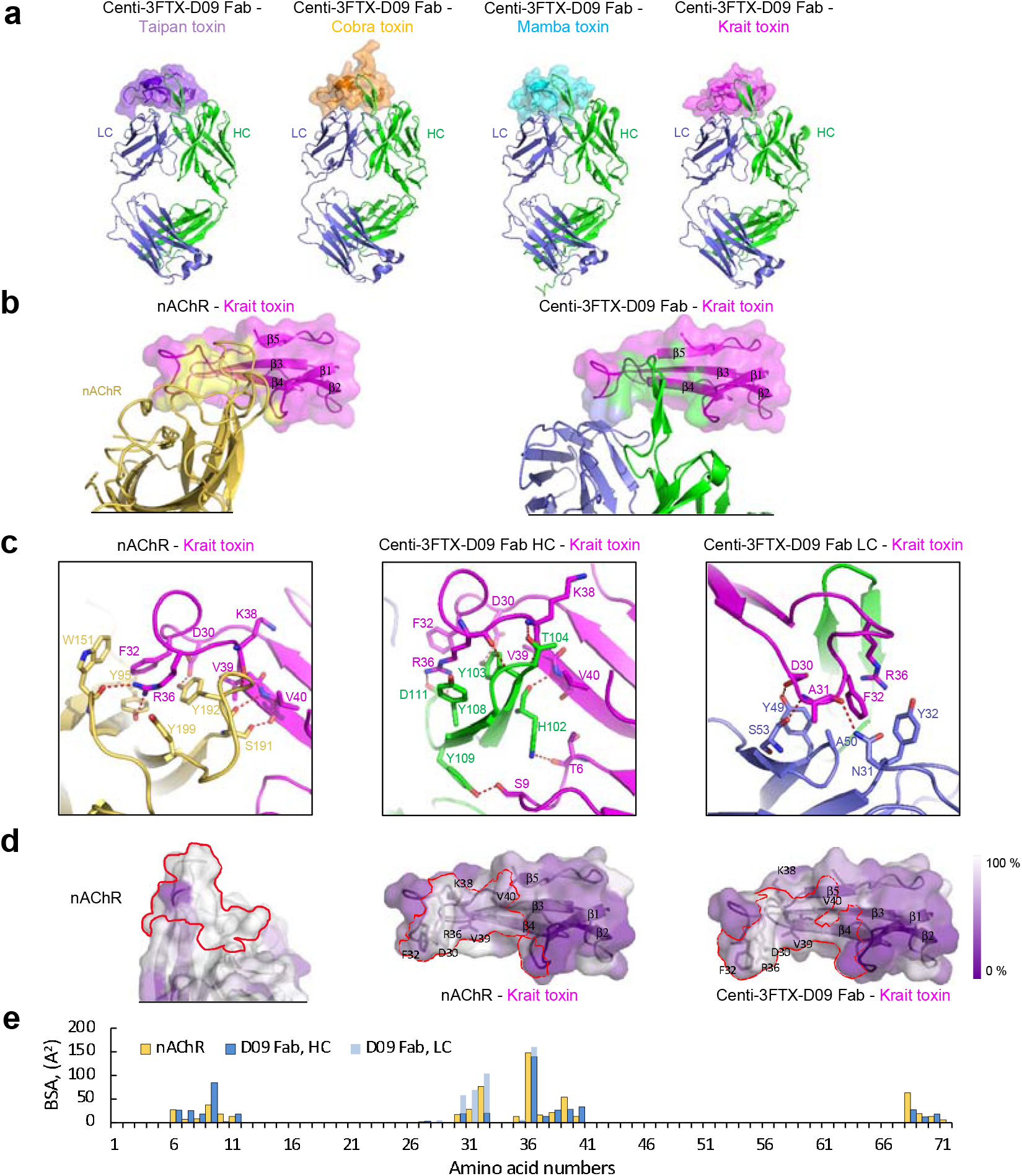
Crystal structures of D09 with long 3FTXs reveal similarity in recognition between antibody and acetylcholine receptor (nAChR). **(a)** Crystal structures of Fab Centi-3FTX-D09 in complex with Taipan, Cobra, Mamba and Krait toxins. **(b)** Toxin recognition of acetylcholine receptor (left) and Centi-3FTX-D09 (right) **(c)** Details, **(d)** Binding surfaces on nAChR and toxin (left); Centi-3FTX-D09 recognized a conserved surface on toxins (right) which closely overlaps with the surface used by toxin to bind acetylcholine receptor (middle). Molecular surfaces colored by sequence conservation, with white indicating 100% conservation. **(e)** Toxin residues involved in binding of Centi-3FTX-D09 and nAChR receptor. Both Centi-3FTX-D09 and nAChR bound the same toxin residues.

Comparison with the previously determined structure of the Krait long 3FTX with the nicotinic acetylcholine receptor^29^ revealed the antibody CDR H3 to approximate the position of “loop C” of the acetylcholine receptor, as part of an interface that was in total ∼15% larger (**Fig. 3b** and **Supplementary Tables 5 and 6**). On the toxin, the key Arg36_Toxin_ extends over Phe32_Toxin_, to form critical hydrogen bonds with the phenol of Tyr95_Receptor_ and the backbone carbonyl of W151_Receptor_; on antibody Centi-3FTX-D09, the aliphatic portion of the Arg36_Toxin_ sidechain extends over Phe32_Toxin_, is further sandwiched by Tyr106_HC_, and forms a bridge with Asp111_HC_ (**Fig. 3c**). Meanwhile, Tyr32_LC_ on the light chain was positioned similarly to Trp151_Receptor_ helping to cradle Phe32_Toxin_ and Arg36_Toxin_.

Analysis of the conservation in acetylcholine receptor and toxin revealed the interface to be highly conserved (**Fig. 3e; Fig. S1; Fig. S2; Online Methods:Conservation**). In general, antibody and receptor interacted with similar toxin residues (**Fig. 3e**).

Collectively, these results reveal antibody mimicry of acetylcholine receptor to enable broad recognition of toxin by antibody.

### In-vivo protection by Centi-3FTX-D09 with challenge by recombinant long 3FTX neurotoxin and whole venom

To determine to what degree broad reactivity *in vitro* would translate to protective neutralization *in-vivo*, we performed recombinant long 3FTX neurotoxin and whole venom challenge experiments in C57BL/6 mice. All animal husbandry and experimental protocols were conducted under the guidance of approved IACUC animal use protocol CR-0119, reviewed, approved and assigned by Charles River D Laboratory (CRADL, South San Francisco, CA) IACUC administrator. (**Online Methods:In-Vivo**).

We began by evaluating whether Centi-3FTX-D09 could provide *in-vivo* protection versus challenge of recombinant long 3FTXs of Cape cobra, common krait, coastal taipan, and black mamba. The recombinant long 3FTX from these four snakes, when injected at the pre-determined LD_100_ of 0.5 mg/kg for common krait and 1.0 mg/kg for the other three species, caused 100 percent lethality starting at 60min to 2 hours, with the exception of one mouse surviving the *N. nivea* injection (**Online Methods:In-Vivo**). Pre-mixed injections of recombinant long 3FTX with 30 mg/kg Centi-3FTX-D09 provided durable protection beyond 24 hours in all cases (**Fig. 4a**). Thus, Centi-3FTX-D09 provided broad neutralizing protection from long 3FTXs from diverse elapids.

**Figure 4.**
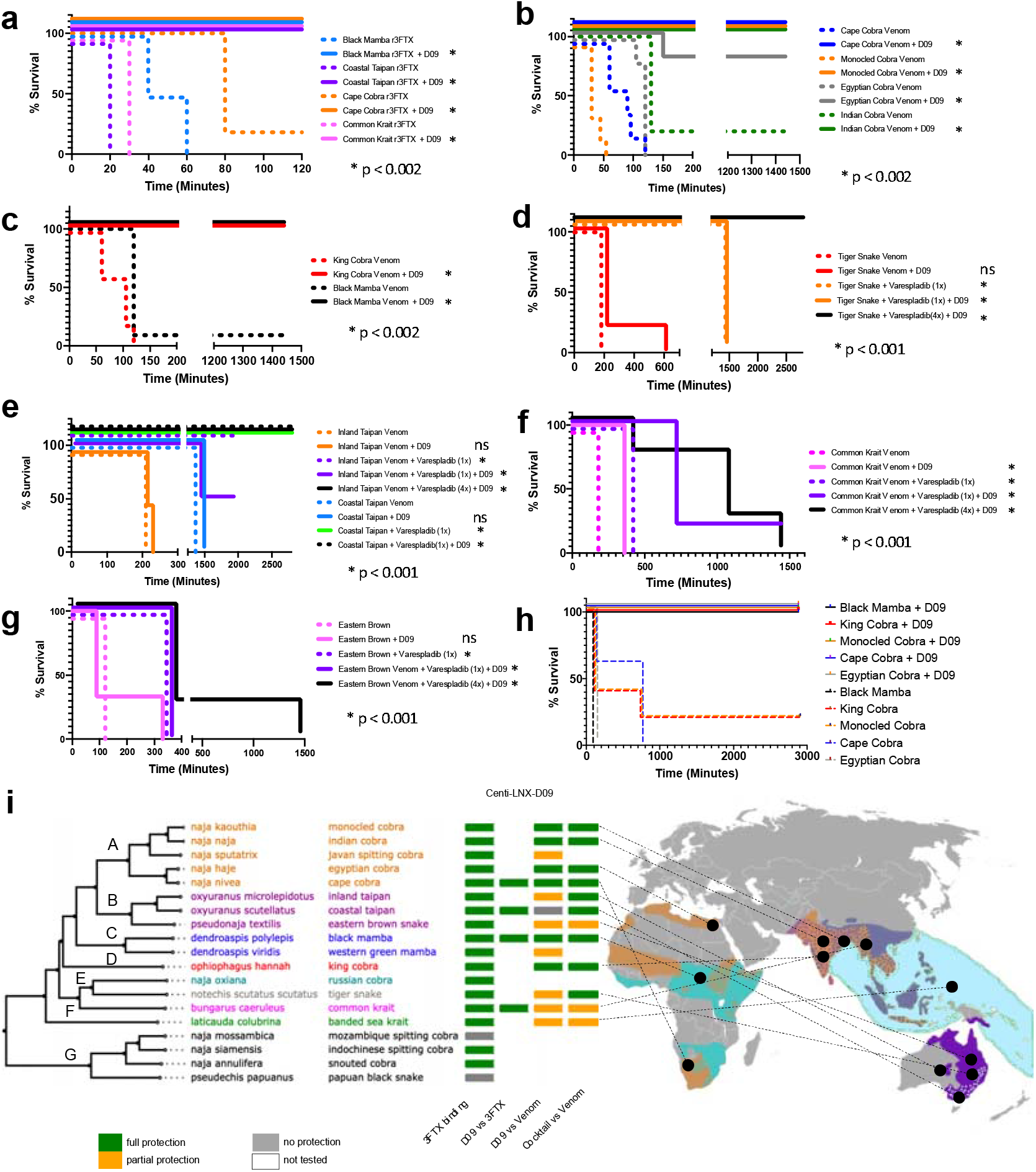
In-vivo protection by D09 with challenge by recombinant long 3FTX neurotoxin and whole venom. Kaplan Meier survival curves for C57BL/6 mice injected **(a-g)** intraperitoneally with venom/antibody premix, or (h) intramuscularly by venom followed by antibody intravenously with 10-minute rescue delay. Centi-3FTX-D09 full protection *in vivo* challenge with **(a)** recombinant long 3FTX from four elapid species (n=10) and whole venom challenge (n=5), including **(b)** four species of cobra, and **(c)** whole venom from two non-cobra elapids. **(d-e)** Full protection for cocktail of Centi-3FTX-D09 and PLA2 inhibitor (varespladib) for **(d)** whole venom from tiger snake, and **(e)** whole venom from two species of taipan. (**f-g**) Partial protection for cocktail of Centi-3FTX-D09 and varespladib for **(f)** common krait and **(g)** eastern brown snake. (h) Centi-3FTX-D09 full rescue protection for mamba, king cobra, and multiple cobras. **(i)** Phylogenetic and geographic breadth of Centi-3FTX-D09 and varespladib *in-vivo* protection to snake venom (Letters above clades map to panels of this figure where venoms were assessed).

Whole venom contains multiple toxic fractions, each of which could contribute to lethality. To deconstruct the relative contribution of the long 3FTX, we first performed whole venom challenge protection studies with Centi-3FTX-D09 as a single agent, on a panel of snakes where the 3FTX was the majority constituent, representing 63-88% of whole venom^16^. As a single agent, Centi-3FTX-D09 provided robust protection from lethal whole-venom challenge for multiple species of cobra (**Fig. 4b**). Complete protection by Centi-3FTX-D09 was observed for *N. nivea* (Cape cobra), *N. kaouthia* (Monocled cobra), and *N. haje* (Egyptian cobra), and for *N. naja* (Indian cobra), 9/10 mice were protected by Centi-3FTX-D09 (**Fig. 4b**). Centi-3FTX-D09 additionally provided complete protection from lethal whole-venom challenge for the non-cobra genera *D. polylepis* (black mamba) and *O. hannah* (King cobra) (**Fig. 4c**). Whole venom protection was observed for both premix IP injections of antibody and venom, as well as rescue studies where animals first received IM injections of venom, and then after a 10-minute delay, received IV injections of Centi-3FTX-D09 (**Fig. 4h**).

After 3FTX, phospholipase A2 (PLA2) is the next most abundant toxin in most elapid venoms. To further deconstruct the contribution of other toxins, we performed whole venom challenge studies with varespladib, a broad-spectrum PLA2 inhibitor, alone and in combination with Centi-3FTX-D09, in a panel of five additional elapid genera.

*Notechis scutatus* (tiger snake) venom, consisting of 74.5% PLA2 and 5.6% 3FTX^16^, was 100% lethal after 2 hours post-injection (**Fig. 4d**). In the mice receiving Centi-3FTX-D09 alone, 1 out of 5 mice survived a total of 10 hours and 4 out of 5 mice survived an additional 30 minutes relative to venom alone. Varespladib alone or in combination with Centi-3FTX-D09 was protective against Tiger Snake lethality for 24 hours, and complete protection was obtained with when varespladib was redosed every 8 hours to account for its short PK.

A similar pattern was observed for taipans. *Oxyuranus microlepidotus* (inland taipan) venom, consisting of 38% PLA2 and 12.2% 3FTX^16^, was 100% lethal in under two hours after injection (**Fig. 4e**). *Oxyuranus scutellatus scutellatus* (coastal taipan) was 100% lethal at 24 hours. The use of Centi-3FTX-D09 marginally extended time to death by a few minutes for venom from both species. Varespladib alone or in combination with Centi-3FTX-D09 was completely protective against a lethal dose of inland and coastal taipan venom (**Fig. 4e**).

*Bungarus caeruleus* (common krait) venom, consisting of 64.5% PLA2 and 19% 3FTX^16^, caused 100% lethality in mice after 3 hours post-injection (**Fig. 4f**). Both Centi-3FTX-D09 and varespladib doubled survival time to 6 and 7 hours, respectively. The combination of Centi-3FTX-D09 and varespladib demonstrated synergistic protective effects, with 4 out of 5 mice surviving 12 hours, and one mouse recovering completely. Redosing with varespladib every 8 hours did not further protect from death.

*Pseudonaja textilis* (eastern brown) venom was 100% lethal after 2 hours after injection (**Fig. 4g**). When Centi-3FTX-D09 was administered with venom, 2 out of 5 mice had extended survival to 6 hours. Varespladib alone or in combination with Centi-3FTX-D09 extended survival of all mice dosed to 6 hours. With repeated redosing of varespladib every 8 hours, two mice survived to 24 hours but then died.

*Laticauda colubrina* (Banded sea krait) venom, consisting of 33% PLA2 and 66% 3FTX^16^, was lethal in 4 out of 5 mice injected 40 minutes after injection. Varespladib alone did not provide protection. Two of 5 mice receiving venom and Centi-3FTX-D09 survived indefinitely (**Extended Data, Fig. S4**). A combination of Varespladib and Centi-3FTX-D09 extended survival of mice to 60-120 minutes, although full protection was not observed.

In the studies discussed so far, we used an envenomation model of intraperitoneal (IP) injections of venom or a premix of venom and Centi-3FTX-D09 antibody. This is of course different from a natural snakebite, where the venom is injected first, typically intramuscularly (IM), and the antivenom is provided afterwards, typically intravenously (IV). Due to the relatively high affinity of interaction between long 3FTX neurotoxins and their nicotinic acetylcholine receptor target, it was important to confirm that Centi-3FTX-D09 would still provide protection when injected following a delay into a mouse already previously envenomed. Thus we repeated the whole venom protection studies of black mamba, king cobra, and multiple cobra species in a rescue model, where venom was injected first IM, and following a 10-minute delay, Centi-3FTX-D09 was injected IV (**Fig. 4h**). In this rescue model, we continued to see complete protection by Centi-3FTX-D09, confirming that the antibody had sufficient affinity to out-compete the nicotinic acetylcholine receptor for the long 3FTX target. This result was consistent with veterinary and human envenomation clinical reports, where antivenom administered after onset of ataxia by neurotoxic snakes has the capacity to rapidly reverse ataxia and rescue the subject.

In a 3FTX phylogeny of the closest 3FTX homolog from each representative elapid species to *N. naja* long 3FTX, Centi-3FTX-D09 provided complete *in-vivo* protection from venom from 6 species spanning 3 genera in clades A (*Naja*), C (*Dendroaspis*), and D (*Ophiophagus*), and partial protection to members of clades B (*Oxyuranus/Pseudonaja*), E (*Notechis/Bungarus/N. oxiana*), and F (*Laticauda*) (**Fig. 4i, Extended Data, Fig. S1**). When combined with varespladib, complete protection was extended to members of clades A-E, spanning Africa, Asia, and Oceana, and partial protection for clade F (oceans). Protection for three species (banded sea krait, eastern brown, and common krait) remained partial for the cocktail at the doses of venom and cocktail evaluated here. Group G (*Pseudoechis/Naja*, particularly spitting cobras), the only clade exhibiting some venoms non-reactive to Centi-3FTX-D09, was not evaluated *in-vivo*.

## Discussion

Here we report Centi-3FTX-D09, a broadly neutralizing anti-long 3FTX antibody capable of binding and neutralizing long 3FTXs from diverse genera of snake venom. A broadly neutralizing antibody against the long 3FTX neurotoxins is of interest, as no small molecule inhibitors for neurotoxins have been identified, unlike the small molecule inhibitors reported for PLA2, SVMP, and SVSP toxins^30^.

As a single agent, Centi-3FTX-D09 provided complete protection from lethal challenge by whole venom from four cobra species, black mamba, and king cobra venom, including full protection in a 10-minute treatment delay rescue model. Centi-3FTX-D09 also provided partial protection against tiger snake, eastern brown, and common krait. Marginal partial protection was also observed for Javan spitting cobra, green mamba, and banded sea krait. Given the diversity of genera recognized, it is likely that Centi-3FTX-D09 could provide protection against venom neurotoxicity for additional members of the 300 venomous species of *elapidae*.

When combined with a broad-spectrum inhibitor of PLA2 (varespladib), we observe extended broad protection against whole venom from a representative panel of 11 snake species, including 8 genera of elapidae, suggesting that 3FTX and PLA2 are the two dominant toxins in the majority of these elapid venoms, although at least one additional toxin continues to drive lethality in a subset of these species, at the doses tested. We observed synergistic protection in some venoms, as well as some venoms that were unilaterally protected by Centi-3FTX-D09 or varespladib alone. Given diversity of genera recognized, it is possible that this cocktail could provide complete protection against whole venom for additional elapids.

In some challenges, we observed delayed death of all animals at 6, 12, or 24 hours following the use of Centi-3FTX-D09 and varespladib as a cocktail. When providing the cocktail followed by redosing of varespladib at 8-hour intervals, full protection was achieved in some additional venoms, confirming that the short (5.5 hour) t_1/2_ of a single injection of *varespladib* was the source of delayed death^31^. In future studies, replacing varespladib with a broadly neutralizing anti-PLA2 antibody may be a path to extend protection indefinitely with a single injection.

We emphasize that these *in-vivo* protection models have limitations. While we observe broad protection across 8 genera at the dose of venom tested, it is known that a live envenomation event may inject more venom than the controlled venom doses used here, and that at such higher doses, additional venom components may increase to a concentration so as to contribute to lethality in a way that is not detected here. Therefore, we interpret the results of these models as a means of identifying the most dominant lethal components of venom, and systematically deconstruct the relative contributions of toxins to lethality by neutralizing individual components and sets of the most toxic components. In future studies, we may choose to increase the dose of venom in fully-protected animals with the current cocktail, in order to uncover any further key toxins to target. We anticipate that it may not be necessary to neutralize all toxins in venom in order to prevent mortality and morbidity from snakebite in humans.

These antibodies were isolated from a hyper-immune individual with a history of heterosubtypic venom immunization exposures that resulted in the generation of potent and broadly cross-neutralizing antibodies against snake venom homologous toxins. From structural and homology analysis of the resulting antibodies, it is evident that the broadly neutralizing antibodies function by exploiting the conserved active sites in this toxin family that have been unable to vary due to a requirement over evolutionary history of being able to target conserved receptor sites in a wide array of vertebrates. As active sites and binding sites are more conserved as a general property of functional proteins, we can anticipate that similar broadly neutralizing antibodies may be able to be recovered for other snake toxin families.

## Supporting information

Ethics statement

## Acknowledgements

We thank Tim Friede for his sample donation and detailed knowledge of herpetology, and members of the Virology Laboratory, Vaccine Research Center, for comments and discussions. Funding was provided in part by the Intramural Research Program of the Vaccine Research Center, National Institute of Allergy and Infectious Diseases, National Institutes of Health, and National Institutes of Health Small Business Innovation Research. Use of sector 22 (Southeast Region Collaborative Access team) at the Advanced Photon Source was supported by the US Department of Energy, Basic Energy Sciences, Office of Science, under contract number W-31-109-Eng-38.

## List of Online Material

Ethics Statement

Online Methods

Extended Data Figure S1. 3FTX conservation.

Extended Data Figure S2. Nicotinic acetylcholine receptor (nAChR) conservation.

Extended Data Figure S3. Centi-3FTX-D09 – recombinant monovalent SPR kinetics.

Extended Data Figure S4. Additional in-vivo whole venom protection studies

Supplementary Table 1. Multiplex HTS+Display adaptor primer sets

Supplementary Table 2. ELISA screening of broadly-reactive antibody scFv clones.

Supplementary Table 3. Identification of immunoprecipitated proteins with mass spectrometry.

Supplementary Table 4. X-ray data collection and refinement statistics.

Supplementary Table 5. Antibody D09-3FTX interface analysis.

Supplementary Table 6. Acetylcholine receptor-Krait 3FTX interface details.

## Figure Legends

**Extended Data Figure S1. 3FTX conservation. (a)** Amino acid sequence alignment of long 3FTX neurotoxins from diverse snake venoms. A single representative 3FTX sequence was obtained for each species. The sequence was chosen as the closest sequence to N. nivea long 3FTX by amino acid identity. In four cases where no long 3FTX was found in UniProt for the species, short 3FTX homolog was instead aligned (*Pseudechis papuanus, Naja siamensis, Naja mossambica, Naja annulifera*). Alignment positional amino acid similarity conservation indicated in blue. **(b)** Pairwise amino acid percent identity (%ID) between long neurotoxin 3FTXs. Yellow=lower %ID, green=higher %ID.

**Extended Data Figure S2. Nicotinic acetylcholine receptor (nAChR) conservation**. A diverse set of representative mammals, marsupials, avians, reptiles, amphibians, and fish were analyzed for conservation of the nAChR receptor (subset of 49 shown above). **(a)** Amino acid sequence alignment of representative nAChRs, cropped to 3FTX interaction domain. Alignment positional amino acid similarity conservation indicated in blue. **(b)** Neighbor-joining tree of representative nAChRs.

**Extended Data Figure S3. Centi-3FTX-D09 binding kinetics. (a)** Biacore Surface Plasmon Resonance Sensograms and residuals at 25C and 37C for monovalent his-avi tagged recombinant “mamba” alpha-elapitotoxin-Dpp2a (3L24_DENPO) from Dendroaspis polylepis (black mamba), “taipan” Long neurotoxin 1 (3L21_OXYSC) from Oxyuranus scutellatus scutellatus (coastal taipan), and “cobra” alpha-cobratoxin/long neurotoxin 1 (3L21_NAJNI) from Naja nivea (Cape cobra), 3FTX interaction with Centi-3FTX-D09. **(b)** Binding kinetic parameter summary tables, including Centi-3FTX-D09 and D09 lineage-related clone Centi-3FTX-B11.

**Extended Data Figure S4**. Additional in-vivo whole venom protection studies. Kaplan Meier survival curves for C57BL/6 mice injected intraperitoneally. Treatments including Centi-3FTX-D09 (30mg/kg) were premixed with venom prior to injection. Treatments with varespladib were pre-treated with a separate injection immediately prior to venom IP injection. For (a) banded sea krait venom, non-statistically significant improvements in survival were observed for either varespladib or Centi-3FTX-D09 alone. In combination, survival was extended from 40 min to 60-120 minutes, but full protection was not observed. For (b) green mamba and Javan spitting cobra, marginal non-statisically significant improvements in survival were observed with Centi-3FTX-D09.

## Notes

### Competing Interest Statement

The authors have declared no competing interest.

### Summary of Updates

Manuscript has been updated to include additional in vivo rescue model data, clarify elements of methodology, and to amend minor textual errors.

