## Supplemental data for "Venom protection by broadly neutralizing antibody from a snakebite subject"

**This PDF file includes:**

Extended Data Figs. S1 to S4

Supplementary Tables S1 to S6

Extended Data Figure S1. 3FTX conservation. (a) Amino acid sequence alignment of long 3FTX neurotoxins from diverse snake venoms. A single representative 3FTX sequence was obtained for each species. The sequence was chosen as the closest sequence to N. nivea long 3FTX by amino acid identity. In four cases where no long 3FTX was found in UniProt for the species, short 3FTX homolog was instead aligned (*Pseudechis papuanus, Naja siamensis, Naja mossambica, Naja annulifera*). Alignment positional amino acid similarity conservation indicated in blue. (b) Pairwise amino acid percent identity (%ID) between long neurotoxin 3FTXs. Yellow=lower %ID, green=higher %ID.

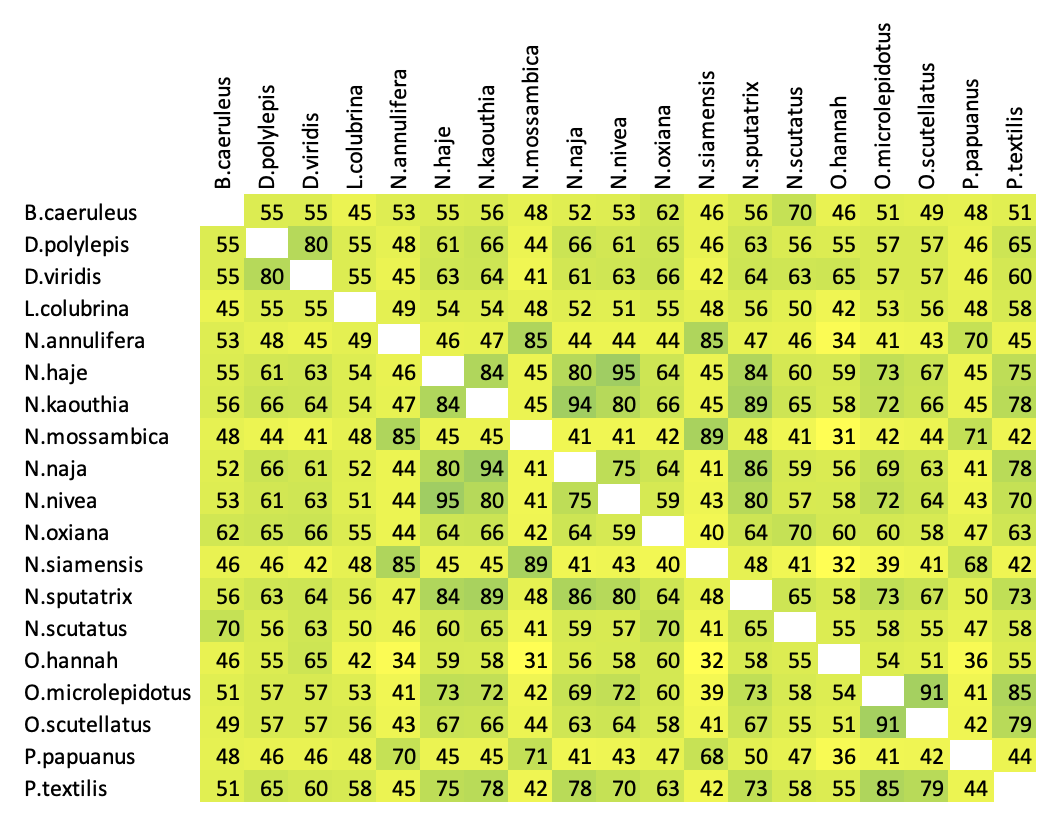

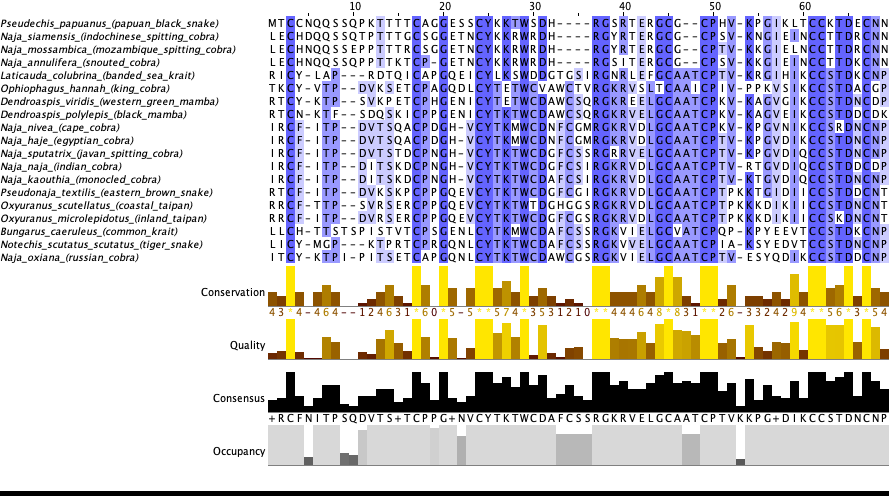

**a**

**b**

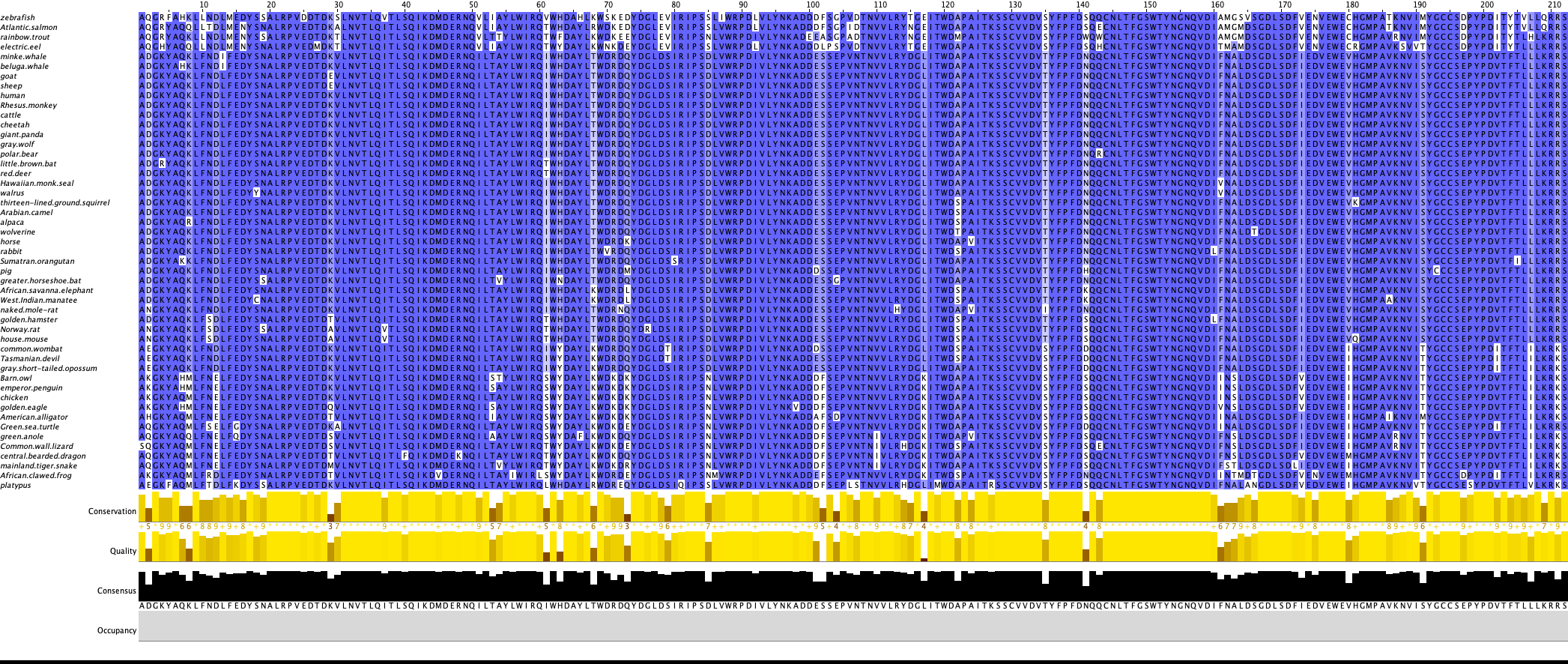

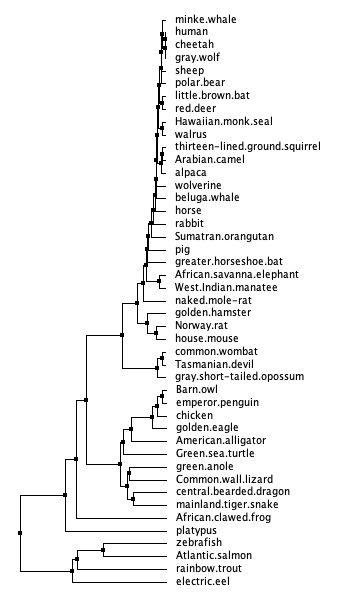

**a**

**b**

**Extended Data Figure S2.** Nicotinic acetylcholine receptor (nAChR) conservation. A diverse set of representative mammals, marsupials, avians, reptiles, amphibians, and fish were analyzed for conservation of the nAChR receptor (subset of 49 shown above). (**a**) Amino acid sequence alignment of representative nAChRs, cropped to 3FTX interaction domain. Alignment positional amino acid similarity conservation indicated in blue. (**b**) Neighbor-joining tree of representative nAChRs.

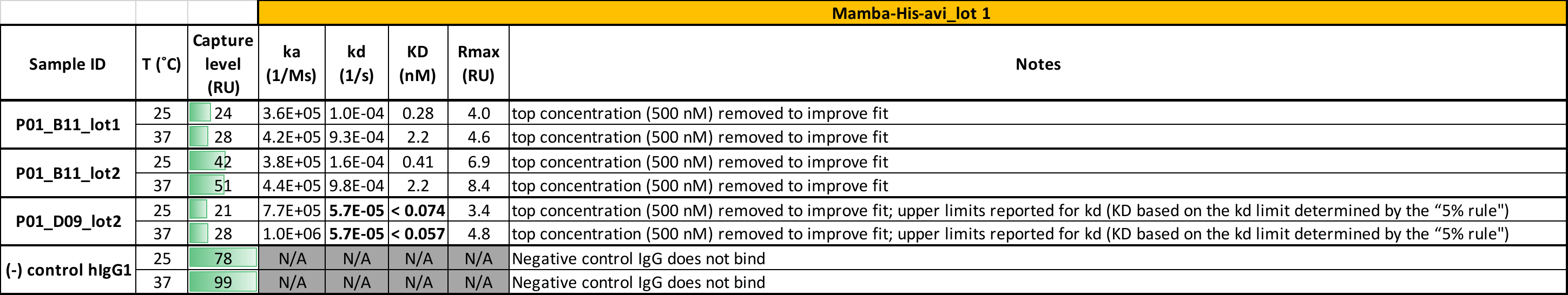

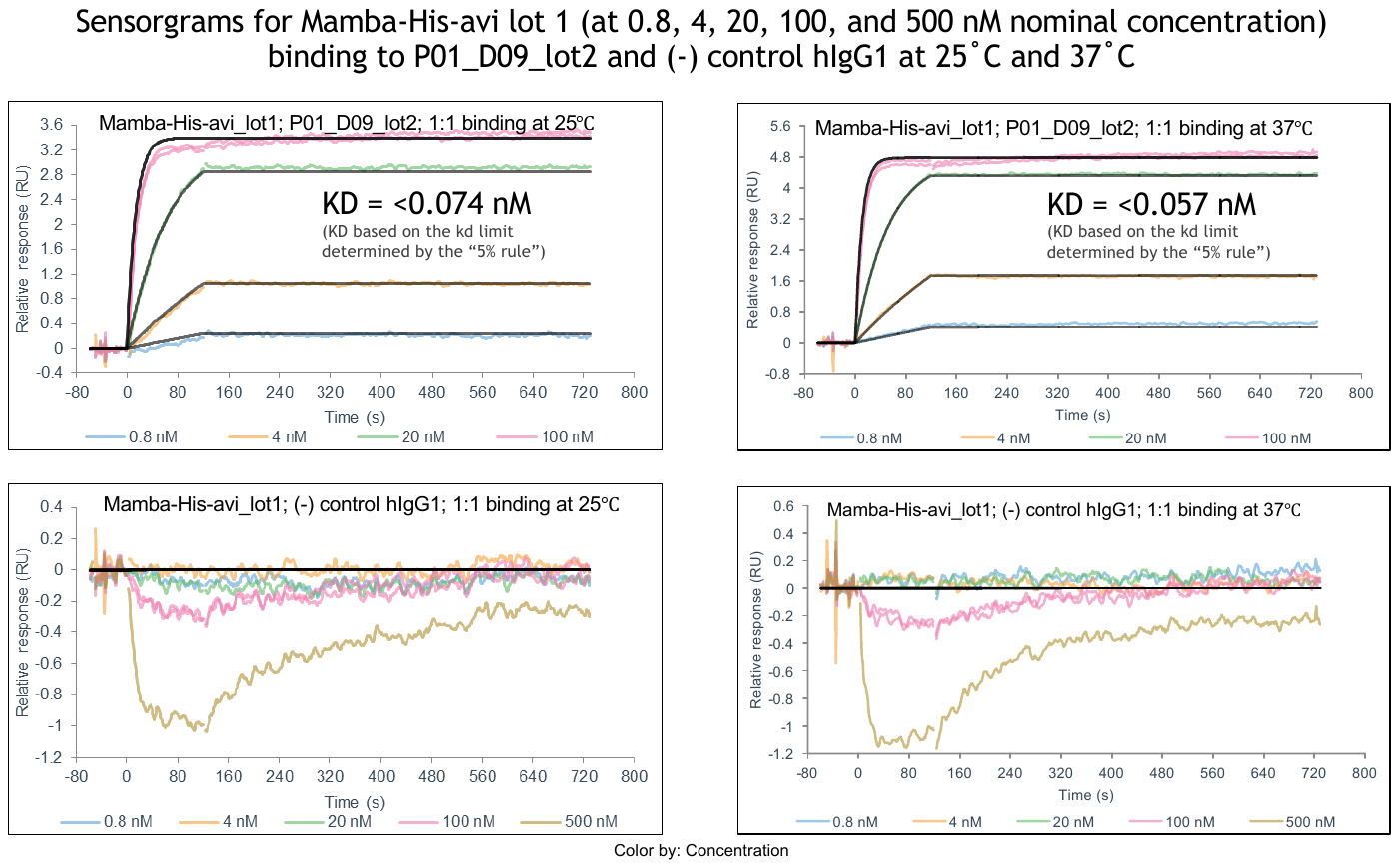

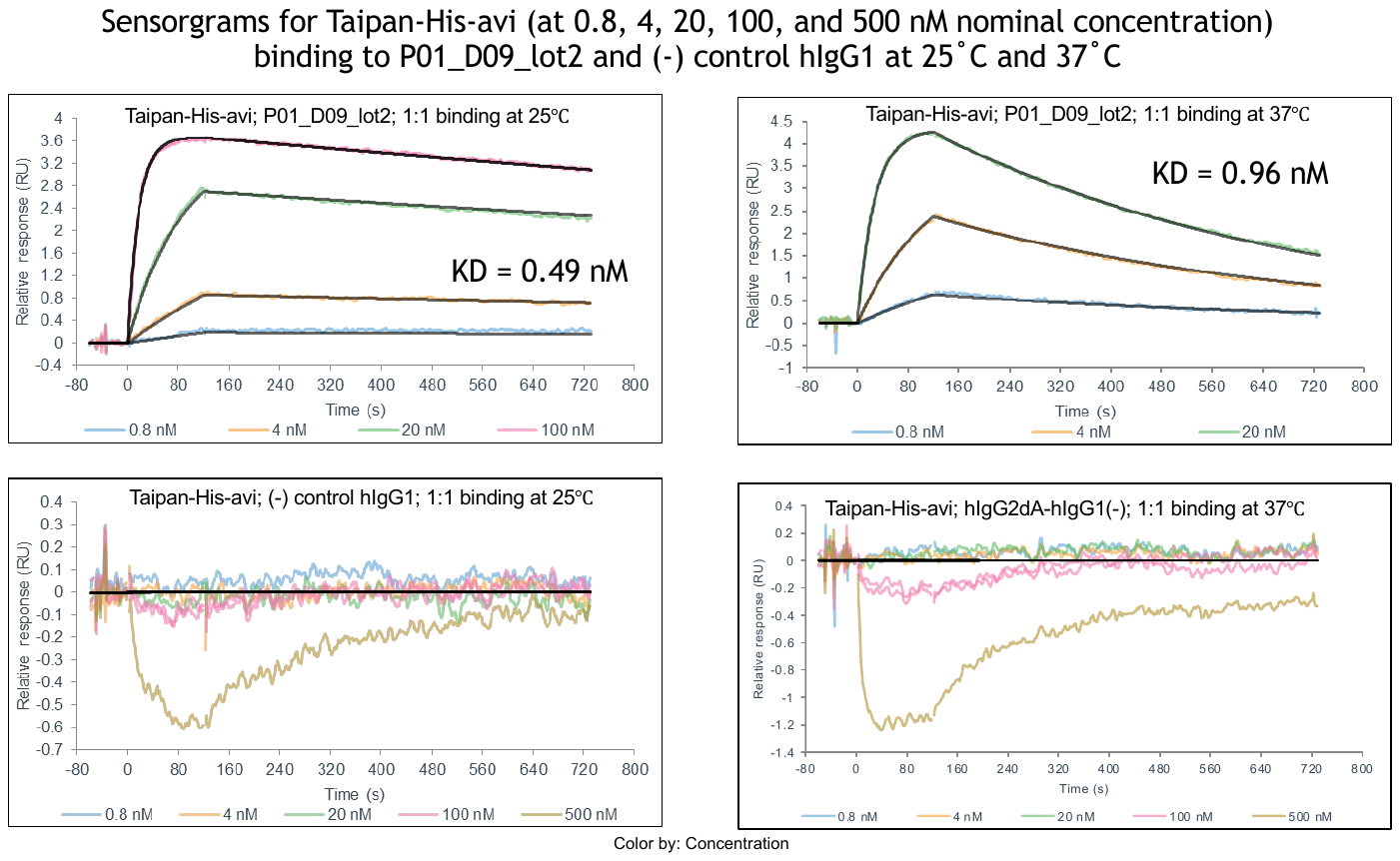

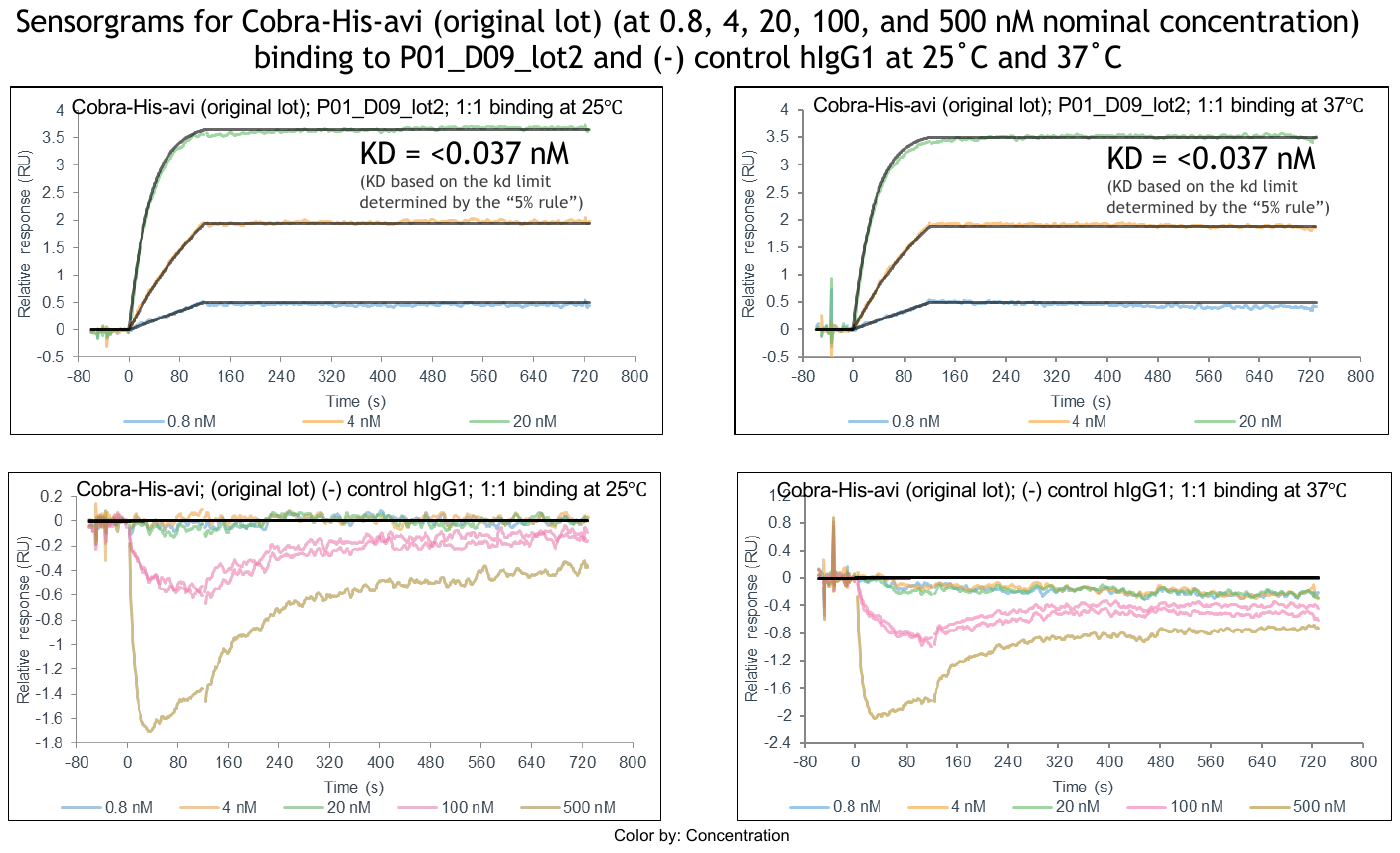

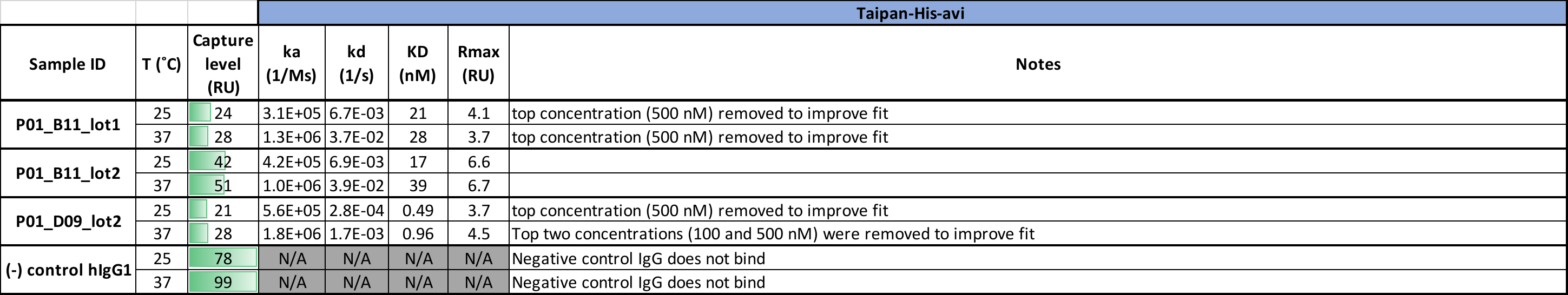

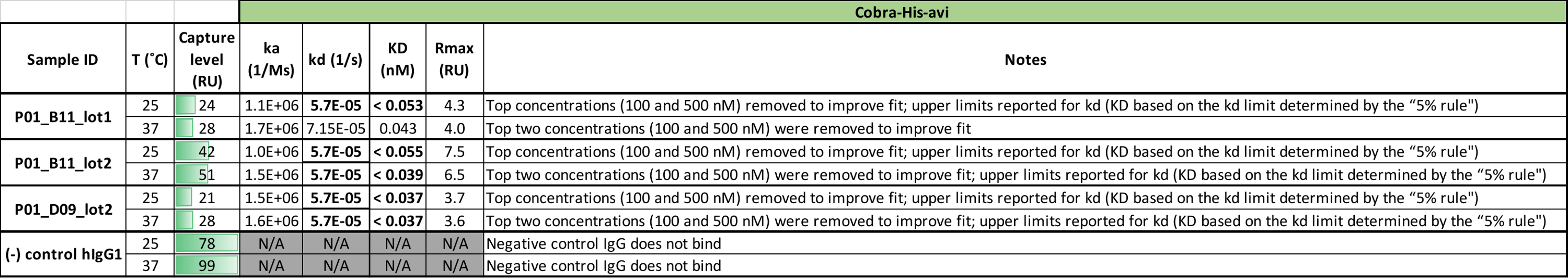

**a**

**b**

**Extended Data Figure S3. Centi-3FTX-D09 binding kinetics.** (**a**) Biacore Surface Plasmon Resonance Sensograms and residuals at 25C and 37C for monovalent his-avi tagged recombinant “mamba” elapitotoxin-Dpp2a (3L24_DENPO) from Dendroaspis polylepis (black mamba), “taipan” Long neurotoxin 1 (3L21_OXYSC) from Oxyuranus scutellatus scutellatus (coastal taipan), and “cobra” alpha-cobratoxin/long neurotoxin 1 (3L21_NAJNI) from Naja nivea (Cape cobra), 3FTX interaction with Centi-3FTX-D09. (**b**) Binding kinetic parameter summary tables, including Centi-3FTX-D09 and D09 lineage-related clone Centi-3FTX-B11.

a

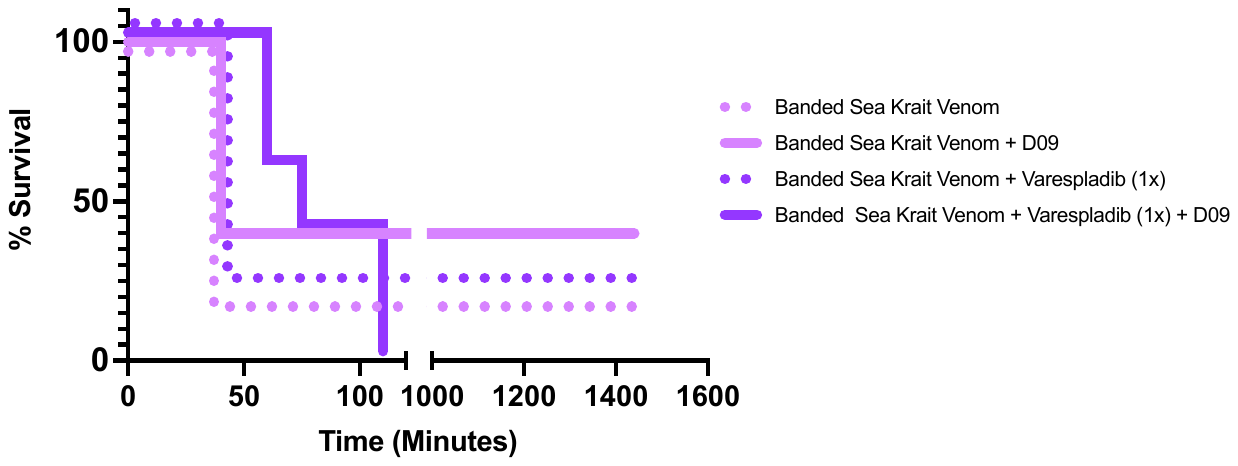

b

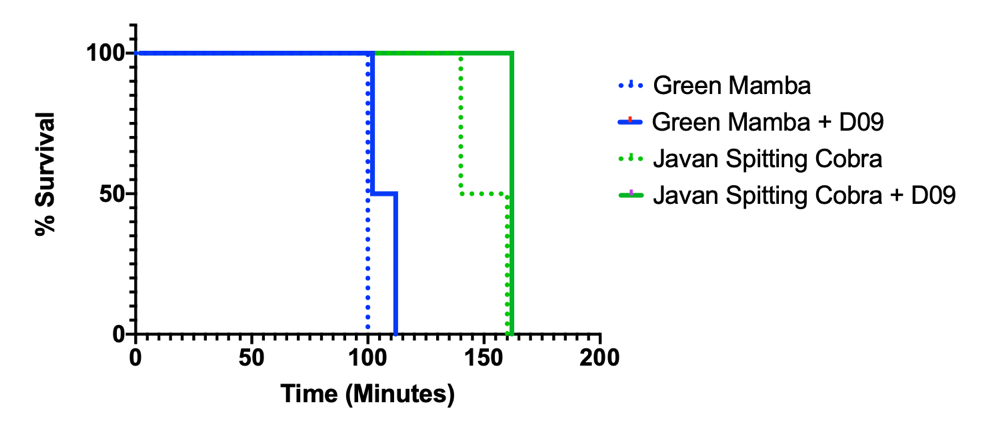

Extended Data, Fig. S4. Additional in-vivo whole venom protection studies. Kaplan Meier survival curves for C57BL/6 mice injected intraperitoneally. Treatments including Centi-3FTX-D09 (30mg/kg) were premixed with venom prior to injection. Treatments with varespladib were pre-treated with a separate injection immediately prior to venom IP injection. For (a) banded sea krait venom, non-statistically significant improvements in survival were observed for either varespladib or Centi-3FTX-D09 alone. In combination, survival was extended from 40 min to 60-120 minutes, but full protection was not observed. For (b) green mamba and Javan spitting cobra, marginal non-statisically significant improvements in survival were observed with Centi-3FTX-D09.

Supplementary Table 1.

Multiplex, dephasing, cloning restriction site embedded, MiSeq flapped-end adapted primer sets for generation of VH, VK, VL amplicons for both high-throughput sequencing and phage display.

| DB-NGS-VK1-NsiI-R2-md01 | GGCATTCCTGCTGAACCGCTCTTCCGATCTNNNNNNTGCGTAGCatgcatccGACATCCAGATGACCCAGTCTCC |
| --- | --- |
| DB-NGS-VK2a-NsiI-R2-md01 | GGCATTCCTGCTGAACCGCTCTTCCGATCTNNNNNNTGCGTAGCatgcatccGATGTTGTGATGACTCAGTCTCC |
| DB-NGS-VK2b-NsiI-R2-md01 | GGCATTCCTGCTGAACCGCTCTTCCGATCTNNNNNNTGCGTAGCatgcatccGATATTGTGATGACCCAGATCCC |
| DB-NGS-VK3-NsiI-R2-md01 | GGCATTCCTGCTGAACCGCTCTTCCGATCTNNNNNNTGCGTAGCatgcatccGAAATTGTGTTGACGCAGTCTCC |
| DB-NGS-VK4-NsiI-R2-md01 | GGCATTCCTGCTGAACCGCTCTTCCGATCTNNNNNNTGCGTAGCatgcatccGACATCGTGATGACCCAGTCTCC |
| DB-NGS-VK5-NsiI-R2-md01 | GGCATTCCTGCTGAACCGCTCTTCCGATCTNNNNNNTGCGTAGCatgcatccGAAACGACACTCACGCAGTCTCC |
| DB-NGS-VK6-NsiI-R2-md01 | GGCATTCCTGCTGAACCGCTCTTCCGATCTNNNNNNTGCGTAGCatgcatccGAAATTGTGCTGACTCAGTCTCC |
| DB-NGS-Vl1-NsiI-R2-md01 | GGCATTCCTGCTGAACCGCTCTTCCGATCTNNNNNNTGCGTAGCatgcatccCAGTCTGTSBTGACGCAGCCGCC |
| DB-NGS-Vl3-NsiI-R2-md01 | GGCATTCCTGCTGAACCGCTCTTCCGATCTNNNNNNTGCGTAGCatgcatccTCCTATGWGCTGACWCAGCCAC |
| DB-NGS-Vl38-NsiI-R2-md01 | GGCATTCCTGCTGAACCGCTCTTCCGATCTNNNNNNTGCGTAGCatgcatccTCCTATGAGCTGAYRCAGCYACC |
| DB-NGS-Vl4-NsiI-R2-md01 | GGCATTCCTGCTGAACCGCTCTTCCGATCTNNNNNNTGCGTAGCatgcatccCAGCCTGTGCTGACTCARYC |
| DB-NGS-Vl7.8-NsiI-R2-md01 | GGCATTCCTGCTGAACCGCTCTTCCGATCTNNNNNNTGCGTAGCatgcatccCAGDCTGTGGTGACYCAGGAGCC |
| DB-NGS-Vl9-NsiI-R2-md01 | GGCATTCCTGCTGAACCGCTCTTCCGATCTNNNNNNTGCGTAGCatgcatccCAGCCWGKGCTGACTCAGCCMCC |
| DB-NGS-Vl11-NsiI-R2-md01 | GGCATTCCTGCTGAACCGCTCTTCCGATCTNNNNNNTGCGTAGCatgcatccTCCTCTGAGCTGASTCAGGASCC |
| DB-NGS-Vl13-NsiI-R2-md01 | GGCATTCCTGCTGAACCGCTCTTCCGATCTNNNNNNTGCGTAGCatgcatccCAGTCTGYYCTGAYTCAGCCT |
| DB-NGS-Vl15-NsiI-R2-md01 | GGCATTCCTGCTGAACCGCTCTTCCGATCTNNNNNNTGCGTAGCatgcatccAATTTTATGCTGACTCAGCCCC |
| DB-NGS-Jk1-NotI-R1-md01 | ACACTCTTTCCCTACACGACGCTCTTCCGATCTNNNNNNTGCGTAGCGCGGCCGCACGTTTGATTTCCACCTTGGTCCC |
| DB-NGS-Jk2-NotI-R1-md01 | ACACTCTTTCCCTACACGACGCTCTTCCGATCTNNNNNNTGCGTAGCGCGGCCGCACGTTTGATCTCCAGCTTGGTCCC |
| DB-NGS-Jk3-NotI-R1-md01 | ACACTCTTTCCCTACACGACGCTCTTCCGATCTNNNNNNTGCGTAGCGCGGCCGCACGTTTGATATCCACTTTGGTCCC |
| DB-NGS-Jk4-NotI-R1-md01 | ACACTCTTTCCCTACACGACGCTCTTCCGATCTNNNNNNTGCGTAGCGCGGCCGCACGTTTGATCTCCACCTTGGTCCC |
| DB-NGS-Jk5-NotI-R1-md01 | ACACTCTTTCCCTACACGACGCTCTTCCGATCTNNNNNNTGCGTAGCGCGGCCGCACGTTTAATCTCCAGTCGTGTCCC |
| DB-NGS-Jl1-NotI-R1-md01 | ACACTCTTTCCCTACACGACGCTCTTCCGATCTNNNNNNTGCGTAGCGCGGCCGCACCTAGGACGGTGACCTTGGTCCC |
| DB-NGS-Jl2-NotI-R1-md01 | ACACTCTTTCCCTACACGACGCTCTTCCGATCTNNNNNNTGCGTAGCGCGGCCGCACCTAGGACGGTCAGCTTGGTCCC |
| DB-NGS-Jl45-NotI-R1-md01 | ACACTCTTTCCCTACACGACGCTCTTCCGATCTNNNNNNTGCGTAGCGCGGCCGCACCTAAAACGGTGAGCTGGGTCCC |
| DB-NGS-Jh1-BmtI-R1-md01 | ACACTCTTTCCCTACACGACGCTCTTCCGATCTNNNNNNTGCGTAGCGCTAGCTGAGGAGACGGTGACCAGGGTGCC |
| DB-NGS-Jh3-BmtI-R1-md01 | ACACTCTTTCCCTACACGACGCTCTTCCGATCTNNNNNNTGCGTAGCGCTAGCTGAAGAGACGGTGACCATTGTCCC |
| DB-NGS-Jh4.5-BmtI-R1-md01 | ACACTCTTTCCCTACACGACGCTCTTCCGATCTNNNNNNTGCGTAGCGCTAGCTGAGGAGACGGTGACCAGGGTTCC |
| DB-NGS-Jh6-BmtI-R1-md01 | ACACTCTTTCCCTACACGACGCTCTTCCGATCTNNNNNNTGCGTAGCGCTAGCTGAGGAGACGGTGACCGTGGTCCC |
| DB-NGS-Vh1-NcoI-R2-md14 | GGCATTCCTGCTGAACCGCTCTTCCGATCTNNNNACTGAGTACCATGGGCCAGGTGCAGCTGGTGCAGTCTGG |
| DB-NGS-Vh2-NcoI-R2-md14 | GGCATTCCTGCTGAACCGCTCTTCCGATCTNNNNACTGAGTACCATGGGCCAGGTCACCTTGAAGGAGTCTGG |
| DB-NGS-Vh3-NcoI-R2-md14 | GGCATTCCTGCTGAACCGCTCTTCCGATCTNNNNACTGAGTACCATGGGCGAGGTGCAGCTGGTGGAGTCTGG |
| DB-NGS-Vh4-NcoI-R2-md14 | GGCATTCCTGCTGAACCGCTCTTCCGATCTNNNNACTGAGTACCATGGGCCAGGTGCAGCTGCAGGAGTCGGG |
| DB-NGS-Vh5-NcoI-R2-md14 | GGCATTCCTGCTGAACCGCTCTTCCGATCTNNNNACTGAGTACCATGGGCGAGGTGCAGCTGTTGCAGTCTGC |
| DB-NGS-Vh6-NcoI-R2-md14 | GGCATTCCTGCTGAACCGCTCTTCCGATCTNNNNACTGAGTACCATGGGCCAGGTACAGCTGCAGCAGTCAGG |

Supplementary Table 2.

Broadly-reactive anti-3FTX clone panel. ELISA screening of periplasmic-extract derived soluble myc tagged scFv vs versus recombinant avi-his tagged 3FTXs from his-avi tagged recombinant “mamba” Alpha-elapitotoxin-Dpp2a (3L24_DENPO) from Dendroaspis polylepis (black mamba), “taipan” Long neurotoxin 1 (3L21_OXYSC) from Oxyuranus scutellatus scutellatus (coastal taipan), “krait” Alpha-delta bungarotoxin (3L2A_BUNCE) from Bungarus caeruleus (common krait, and “cobra” alpha-cobratoxin/long neurotoxin 1 (3L21_NAJNI) from Naja nivea (Cape cobra). ELISA screening reported as max OD450 signal, fold over negative control background. Yellow: higher binding signal, Red: lower binding signal, colors relative to within-species ELISA scores. Sequence frameworks and CDR sequences reported. Analysis by VDJFasta (*18*).

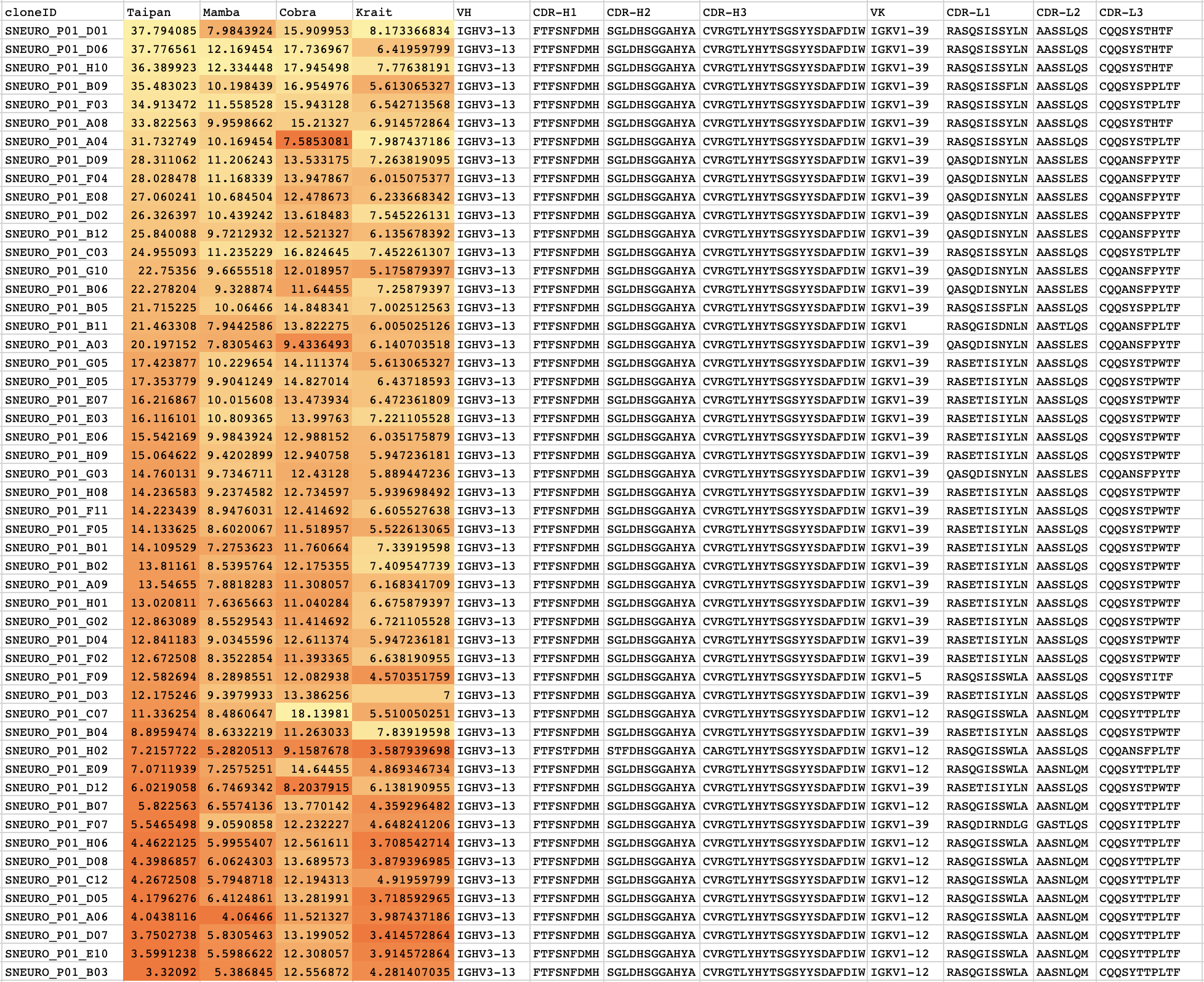

Supplementary Table 3.

Mass spectrometry of Centi-3FTX-D09 -bound proteins from snake venom.

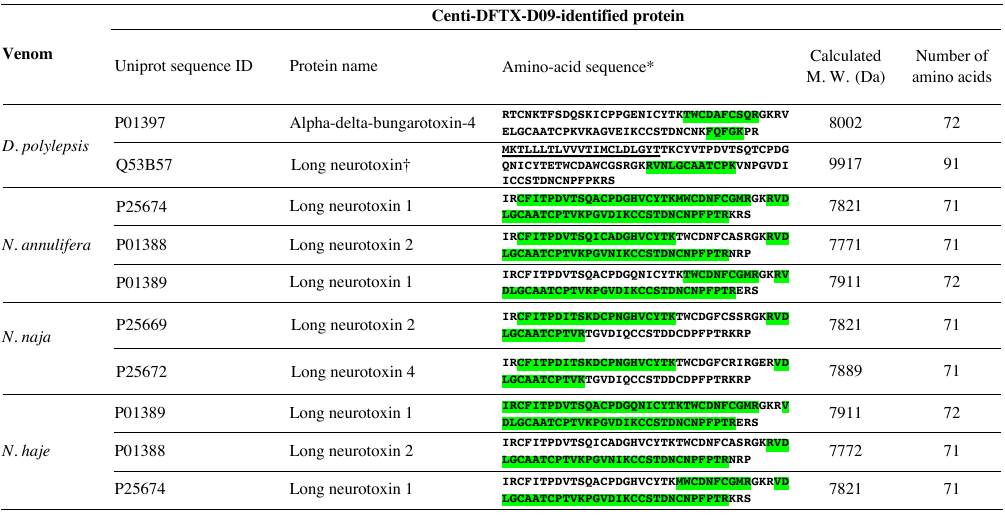

* Sequence segments highlighted in green were identified by mass spectrometry.

† This Uniprot sequence ID contains a signal peptide, which is underlined in the amino-acid sequence.

Supplementary Table 4.

Data collection and refinement statistics.

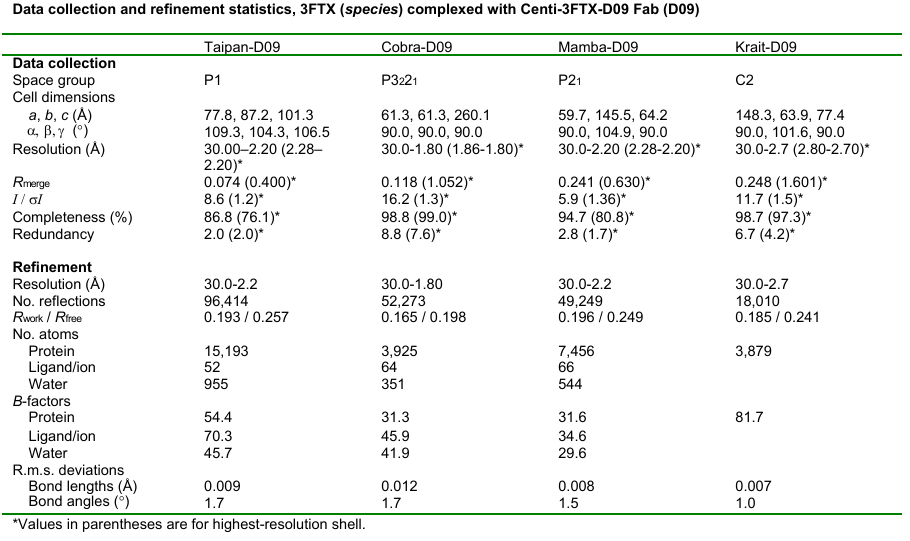

Supplementary Table 5. Antibody
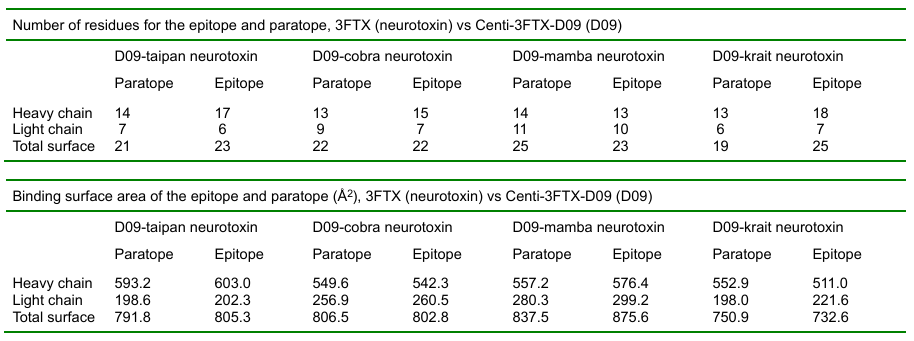
Centi-3FTX-D09 Fab – 3FTX interface overall summary

**Table S5a.** Centi-3FTX-D09 – 3FTX interface overall summary

**Supplementary Table 5b**. Centi-3FTX-D09 Fab – taipan 3FTX neurotoxin interface details

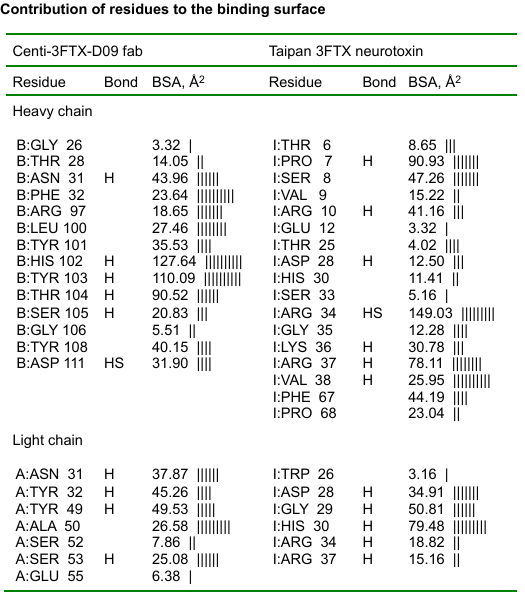

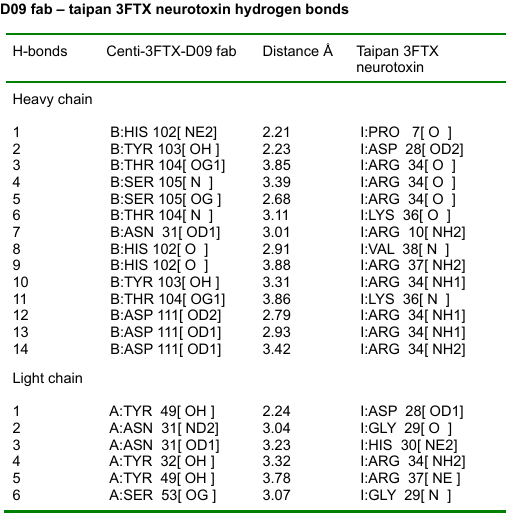

**Supplementary Table 5c**. Centi-3FTX-D09 Fab – cobra 3FTX neurotoxin interface details

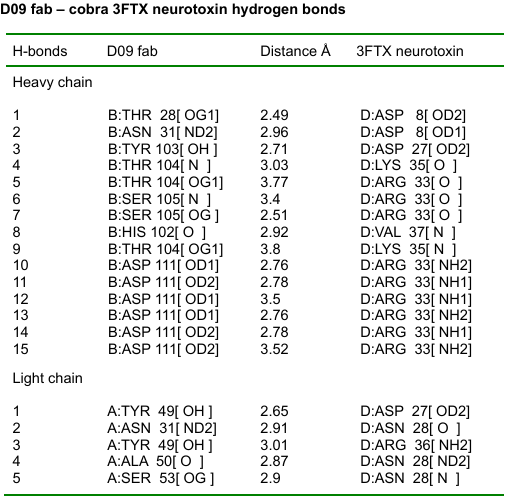

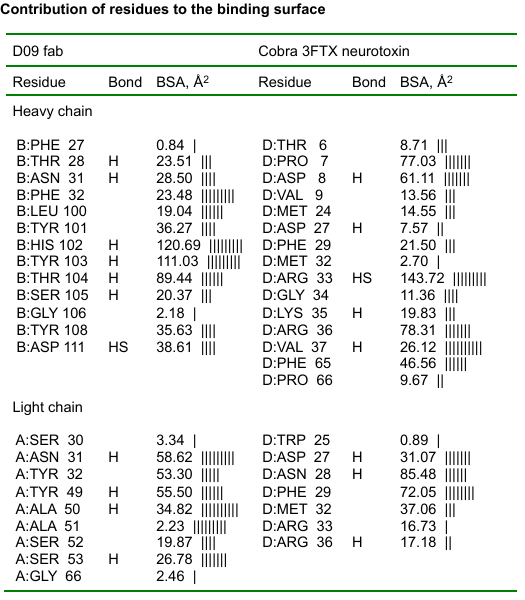

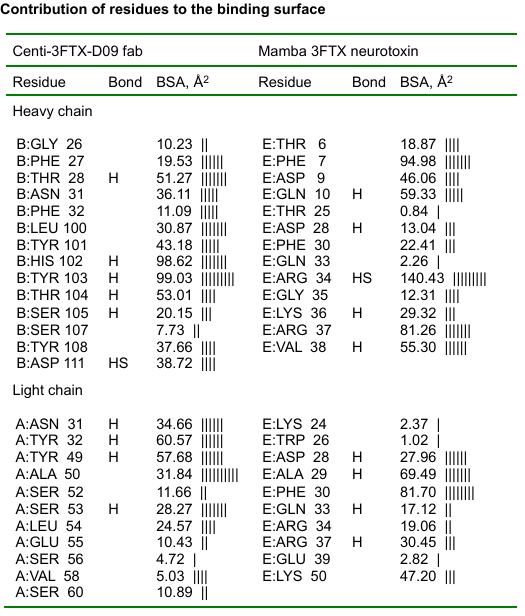

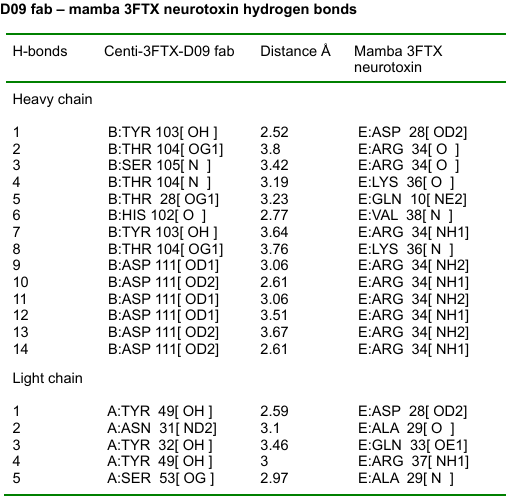
**Supplementary Table 5d**. Centi-3FTX-D09 Fab – mamba 3FTX neurotoxin interface details

**Supplementary Table 5e**. Centi-3FTX-D09 Fab – krait 3FTX neurotoxin interface details

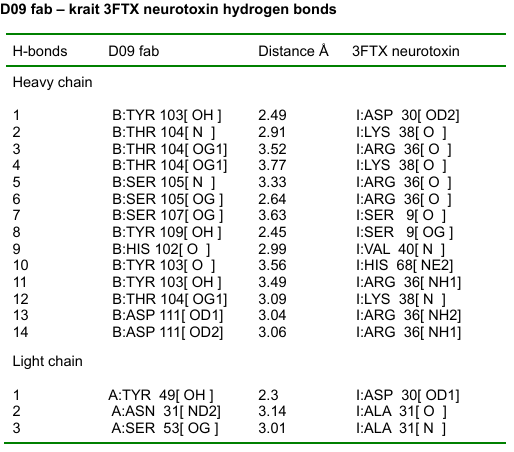

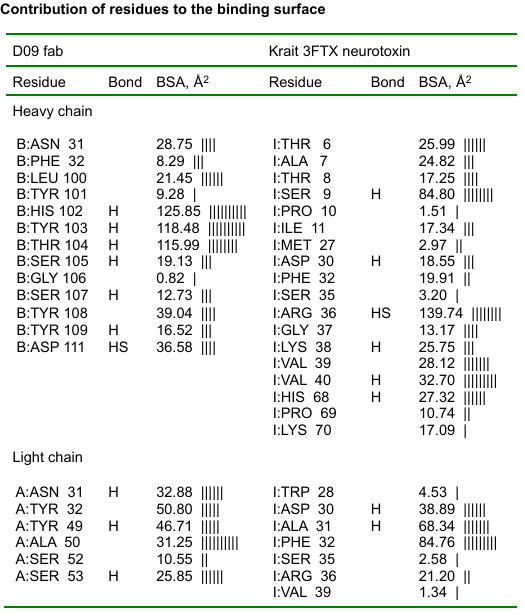

Supplementary Table 6.

Acetylcholine receptor- Krait 3FTX alpha-bungarotoxin interface details.

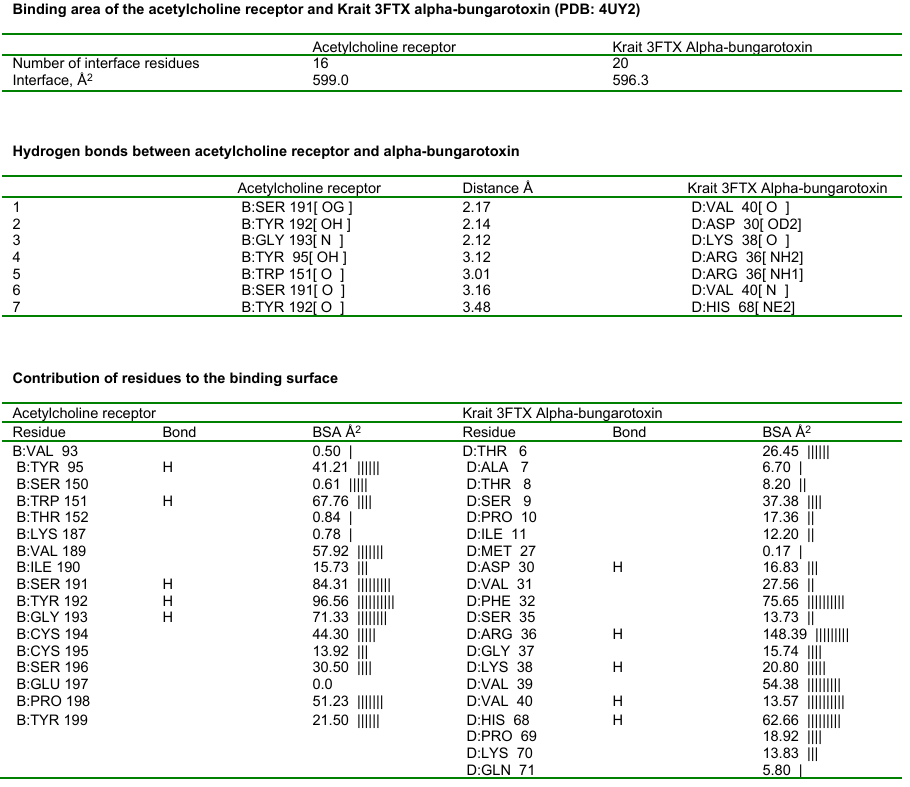
