## Supplementary material for "Venom protection by broadly neutralizing antibody from a snakebite subject": Methods

Methods for

**This PDF file includes:**

Methods

Methods

Plasma & PBMCs

Blood was obtained by **ExamOne** in 4 10ml purple top EDTA anti-coagulant vials. Plasma and cells from each blood draw were isolated by centrifugation at 25°C for 10 minutes at 300xg. Informed consent was obtained from the subject. Western Institutional Review Board (WIRB, now wcgIRB) provided IRB exemption determination #1-1209200-1 for the study of the samples of a single subject (1019 39^th^ Avenue SE Suite 120, Puyallup, CA 98374).

Serum ELISA

One hundred nanograms of each recombinant, C-terminal his-avi tagged long-neurotoxin protein from Alpha-elapitotoxin-Dpp2a (3L24_DENPO) from *Dendroaspis polylepis* (black mamba), alpha-cobratoxin/long neurotoxin 1 (3L21_NAJNI) from *Naja nivea* (Cape cobra), Long neurotoxin 1 (3L21_OXYSC) from *Oxyuranus scutellatus scutellatus* (coastal taipan), and Alpha-delta bungarotoxin (3L2A_BUNCE) from *Bungarus caeruleus* (common krait) were added to microtiter plates (Corning), in phosphate-buffered saline (PBS). After incubation at 4°C overnight and blocking with 3% bovine serum albumin (BSA) (Sigma-Aldrich) in PBS, for 1 hour at 37°C, serially diluted plasma (fivefold, six dilutions, starting from 1:100) in blocking buffer was added to wells and incubated for 1 hour at 37°C. Then, plates were washed three times with 0.05% (v/v) Tween 20 (Sigma-Aldrich) in PBS (PBST). Horseradish peroxidase (HRP)–conjugated donkey anti-human IgG Ab (Jackson Immuno Research Labs) was added to wells and incubated for 1 hour at 37°C. After washing three times with PBST, 2,2′-azino-bis-3-ethylbenzothiazoline- 6-sulfonic (Thermo Fisher) was added to the wells. Absorbance was measured at 415 nm using a microplate spectrophotometer (Multiskan Skyhigh, Thermo Fisher). All experiments were performed in triplicate, and data are presented as the mean ± SD.

Amplicons

PBMC RNA was extracted from cells by RBC lysis followed by RNeasy Mini Kit according to the manufacturer’s protocol (QIAgen). PBMC lysate was homogenized via QIAshredder spin columns (QIAgen) prior to use of RNeasy Mini Spin columns. Following RNA isolation, cDNA was generated using High Capacity cDNA Reverse Transcription Kit (ThermoFisher) following the manufacturer’s protocol. Antibody V-genes (VH, VK, VL) were then amplified from cDNA using custom designed HTS+Display primers to add multiplexed dephasing adapted ends for Illumina MiSeq loading as well as Fv-boundary restriction sites to enable direct digestion and library construction (**Table S1**).

PCR amplification reactions were run with adapted primers at 10 uM using OneTaq Polymerase in GC Buffer (New England Biolabs) at the following cycling conditions: initial denaturation of 2 minutes at 94^o^C, thirty cycles of 94^o^C denaturation for 30 seconds, 53^o^C annealing for 30 seconds, 68^o^C extension for 45 seconds, and a final extension of 68^o^C for 2 minutes. A second Paired End PCR reaction was run to further amplify the resulting MiSeq library as well as add additional bases required for hybridization to the MiSeq flow cell with primers dubbed OptPE1 and OptPE2. This reaction was performed with primers at 10 uM and Phusion Polymerase with GC Buffer (New England Biolabs) at the following cycling conditions: initial denaturation of at 98^o^C for 30 seconds, followed by 12 cycles of 98^o^C denaturation for 8 seconds, 69^o^C annealing for 12 seconds, 72^o^C extension for 10 seconds, and a final extension of 72^o^C for 5 minutes. Each domain was deep sequenced by MiSeq (2x300 v3 kit).

Library

Amplicons were digested and cloned into M13 pIII fusion phage display scFv vector in the VH-VL -pIII orientation, with a 15-mer (G4S1)3 linker bounded by restriction sites, method adapted from previously described methods (*1*, *2*). The library was cloned into two sub-libraries: one utilizing the VK amplicon as light chain and one using the VL amplicon. The VH/VK library was electroporated into ER2738 bacterial phage display electrocompetent cells (Lucigen) at 9.24e8 transformants and the VH/VL library at 1.28e9 transformants for a total library size of 2.21e9 transformants. The resulting library was then rescued using M13K07 Helper Phage (New England Biolabs) at a Multiplicity of Infection (MOI) of 20.

Recombinants

From an analysis of 6706 toxins from 703 snake species in Uniprot database(*3*), Alpha-elapitotoxin-Dpp2a (3L24_DENPO) from *Dendroaspis polylepis* (black mamba), alpha-cobratoxin/long neurotoxin 1 (3L21_NAJNI) from *Naja nivea* (Cape cobra), Long neurotoxin 1 (3L21_OXYSC) from *Oxyuranus scutellatus scutellatus* (coastal taipan), and Alpha-delta bungarotoxin (3L2A_BUNCE) from *Bungarus caeruleus* (common krait), and 21 other toxins were selected for recombinant expression. Toxins were assigned C-terminal tags, including toxin-Fc-avitag fusions and toxin-10his-avitag fusions. Fc-tag and 10his tags were provided to assist in toxin purification, and avitags were provided to enable site-specific biotinylation to magnetic beads for automated soluble phase panning. When induced for expression in HEK293, 21 of the 25 toxins expressed. Toxins were tag-purified (protein G Dynabeads or nickel bead for Fc and 10his tag toxins, respectively), and 15 constructs had strong reactivity to hyperimmune donor serum over healthy control serum by Octet QK. Confirmation of neurotoxic functional activity was performed in-vivo in B6 mouse Maximum Tolerated Dose (MTD) studies, with Alpha-elapitotoxin-Dpp2a from *Dendroaspis polylepis* (black mamba), alpha-cobratoxin/long neurotoxin 1 from *Naja naja* (Indian cobra), Long neurotoxin 1 from *Oxyuranus microlepidotus* (inland taipan), and Alpha-delta bungarotoxin from *Bungarus caeruleus* (common krait) confirmed to be lethal (LD100) at 0.5-1mg/kg.

Panning

Toxins were site-specific biotinylated at C-terminal avitags and quality-controlled for biotinylation (*1*, *2*). The Hyperimmune library (2.21 × 10^9^) was heated for 10 min at 72 °C and de-selected against Protein G Dynabeads^TM^ (Invitrogen), M-280 Streptavidin Dynabeads^TM^ (Invitrogen), Histone from Calf Thymus (Sigma), Human IgG (Sigma) and ssDNA-Biotin NNK from Integrated DNA Technologies and DNA-Biotin NNK from Integrated DNA Technologies. Next, the library was panned against the toxin-captured by M-280 Streptavidin Dynabeads^TM^ using an automated protocol on Kingfisher FLEX (Thermofisher). Selected phages were acid eluted from the beads and neutralized using Tris-HCl pH 7.9 (Teknova). ER2738 cells were infected with the neutralized phage pools at OD_600_ = 0.5 at a 1:10 ratio and after 40 min incubation at 37 °C and 100 rpm, the phage pools were centrifuged and incubated on agar with antibiotic selection overnight at 30 °C. The rescued phages were precipitated by PEG and subjected to three additional rounds of soluble-phase automated panning. PBST/1% BSA buffer and/or PBS/1% BSA was used in the de-selection, washes and selection rounds. Four rounds of Panning were performed on each of the libraries using site-specific avitag biotinylated recombinant long 3FTX as the antigen with reduced concentration in each round (100nM, 20nM, 4nM and, 1nM, respectively) to ensure high selectivity. A different ortholog of long 3FTX was used in each subsequent round to select for breadth: Alpha-elapitotoxin-Dpp2a (3L24_DENPO) from *Dendroaspis polylepis* (black mamba) in round 1, Long neurotoxin 1 (3L21_OXYSC) from *Oxyuranus scutellatus scutellatus* (coastal taipan) in round 2, alpha-cobratoxin/long neurotoxin 1 (3L21_NAJNI) from *Naja nivea* (Cape cobra) in round 3, and Alpha-delta bungarotoxin (3L2A_BUNCE) from *Bungarus caeruleus* (common krait) in round 4.

PPE Screening

Following panning, 184 scFv clones were induced for expression by Isopropyl β-D-1-thiogalactopyranoside (IPTG) via the lac operon incorporated in the M13 pIII library vector from each panning output. Periplasmic extract from these clones were screened for binding to the recombinant toxins from which they were panned by ELISA using pre-coated and pre-BSA blocked 96-well ELISA Plates (Pierce) to capture the toxins at 2 ug/mL.

Informatics

Sanger and HTS sequences were analyzed by VDJFasta2.0 (*4*), utilizing HMMER v3.2 Fv Hidden-Markov Models and NCBI Blast (*5*, *6*) . Figures were rendered with Prism and in R.

The Centi-3FTX-D09 heavy chain has long CDR-H3 and bears evidence of extensive somatic hypermutation. Centi-3FTX-D09 utilized the IGHV3-13 V-gene framework, VDJ-recombination with IGHD3-10 to form a long (19 aa) CDR-H3 loop. Centi-3FTX-D09 had been highly mutated, 77.4% ID to germline with SHM mutational evidence of affinity maturation, with 21 amino acid mutations relative to germline IGHV3-13*01 (93 positions observable: as first 5 positions and j-segment are impacted by primers that can alter amino acid content and therefore masked from analysis, and CDR-H3 VDJ recombination results in non-templated sequence that cannot be inferred, and therefore filtered from analysis). The mutations were concentrated in the CDRs and particularly concentrated in CDR-H2, with 2 mutations in CDR-H1, 9 mutations in CDR-H2, and one mutation in CDR-H3 vernier Kabat 93. Three additional affinity maturation variants of Centi-3FTX-D09 VH were recovered, all showing evidence of extensive SHM. Centi-3FTX-B11 affinity maturation variant, also had broadly neutralizing activity against the target, although not as potent as Centi-3FTX-D09.

The Centi-3FTX-D09 light chain was subjected to diversification mutagenesis analysis, revealing total restriction to IGKV1, strong specific restriction to IGKV1-39, and a strong consensus with polymorphic positions concentrated in CDRs. During immune library generation, all antibody VK domains in the subject's sampled repertoire were associated with each heavy chain. After selection, 118 positive clones were recovered belonging to the Centi-B9 VH CDR-H3 lineage. The clones contained 14 unique light chains. The light chains were entirely restricted to four IGKV1 V-segment family members, with 100 (85%) belonging to IGKV1-39, 13 (11%) to IGKV1-12, 3 (2.5%) IGKV1-5, 1 (0.8%) IGKV1-6, and one (0.8%) unresolvable beyond IGKV1 due to sequence quality. Across the 14 unique light chain variants, 37 positions were observed to tolerate mutations, with only 9 positions with simpson's index greater than 0.5: 4 in CDR-L1, 1 in CDR-L2, 4 in CDR-L3. In the Centi-3FTX-D09 clone, there were 8 positions altered relative to germline (3 L1, 1 L2, 1 FW3, 3 putative L3 positions), although entirely germline IGKV1-39 light chains were also identified to possess broadly reactive binding.

Kinetics

The kinetics and affinities for the interactions of Centi-3FTX-D09 with a-neurotoxins at 25 °C and 37 °C were determined on a Biacore 8K SPR instrument (Cytiva, Marlborough, MA). An anti-human Fc capture chip was prepared by amine-coupling a goat anti-human IgG Fc antibody (catalog No. 2014-01, Southern Biotech, Birmingham, AL) to a Biacore Series S CM4 sensor chip (catalog No. BR100534, Cytiva, Marlborough, MA). The running buffer for the immobilization procedure at 25 °C was HBS supplemented with 0.05% (v/v) Tween-20. The anti-human Fc was diluted to 50 µg/mL into 10 mM sodium acetate pH 4.5, and injected in all flow cells at 20 µL/min for 7 min after activation of the surfaces with a 1:1 (v/v) mixture of 400 mM EDC and 100 mM N-hydroxysuccinimide (NHS) for 7 min at 10 µL/min. Excess reactive esters on the surface of flow cells 1 and 2 were blocked for 7 min at 10 µL/min with 100 mM ethylenediamine in 200 mM borate buffer pH 8.5. Kinetic assays were conducted at 25 °C and 37 °C with HBS supplemented with 0.05% (v/v) Tween-20 and 1 mg/mL BSA as running buffer. The Centi-3FTX-D09 bnAb was diluted to 10 µg/mL with running buffer and captured by its Fc onto the anti-human Fc surface for 2 min at 10 µL/min. His tagged toxins were injected as analyte for 2 min followed by a 15 min dissociation at a flow rate of 30 µL/min. Five analyte concentrations were – 1.2, 3.7, 11.1, 33.3 and 100 nM in addition to a buffer analyte cycle. The anti-human Fc surfaces were regenerated using 3, 60-s injections of 75 mM phosphoric acid at 10 µL/min. Kinetic data were double-referenced (Myszka, 1999), and fit globally to a simple 1:1 Langmuir binding model using the Biacore Insight Evaluation Software (version 3.0). (*7*)

Antibody immunoprecipitation of 3FTX from-venom with mass spectroscopy identification

Lyophilized venom was dissolved in PBS as 50mg/ml solution. 1mg venom solution was mixed with 150ug (150ul) of antibody IgG. 100ul Protein A resin (50 ul settled resin volume, GE Health Sciences) were added to venom-antibody mixture. The mixture was then incubated on a rotating shaker at room temperature for 1 hour. After incubation, the mixture was loaded on an empty Ployprep column (BioRad). The resin in the column was washed twice with 3ml PBS using gravity flow. The resin was resuspended in PBS buffer and transferred to a 1.5ml microtube. Resin was spined down and clarified solution was removed. Bound IgG and toxin proteins were eluted with 50ul SDS-PAGE gel loading buffer (Thermo Fisher Scientific). Proteins bound to the Protein A resin was then separated on SDS-PAGE gel. The bands run around 6 kDa were cut off from the gel and were sent for mass spectrum (LC/MS/MS) analysis at Poochon Scientific (Frederick, Maryland). Protein gel-band samples were digested using trypsin/lysC. The LC/MS/MS analysis was carried out using a Thermo Scientific Orbitrap Exploris 240 Mass Spectrometer and a Thermo Dionex UltiMate 3000 RSLCnano System. The instrument was operated in the data dependent mode to automatically switch between full scan MS and MS/MS acquisition. The MS raw files were analyzed using Proteome Discoverer 2.4 against snake protein sequence database containing 6706 toxin proteins from 774 species downloaded from NCBI and UniProtKB websites (*3*, *5*). The proteins with matched peptides sequences identified from each sample are summarized in **Table S**3.

Conservation

3FTX diversity analysis: Sequences for all 3FTX long neurotoxins were obtained from UniProt SWISS-PROT by both sequence homology to representative PDB ID:1ABT (blastp, e-val: 1e-5) and by sequence annotation (*3*). Sequences were aligned by muscle to structural reference 1ABT, partial sequences were removed, and rendered non-redundant at 99% amino acid identity. Simpson's diversity index D was calculated for each position in the alignment.

nAchR diversity analysis: Two-hundred and fifty (250) putative ortholog amino acid sequences of human reference nAchR were obtained from UniProtKB by blastp (e<1-e5) and aligned by Muscle to human nAchR crystallographic reference structure of the ligand-binding domain PDBID:4UY2 (*3*, *5*). Common names for each species were obtained by lookup against UniProtKB-provided species binomial nomenclature in NCBI taxonomy: mammals, avians, amphibians, reptiles, fish and sharks were identified. Sequences were curated to remove duplicates or potential paralogs per species and sequence fragments, leaving 246 sequences in the dataset. UniprotKD annotations were reviewed to confirm sequence selection criteria successfully selected putative orthologs. Sequences were cropped to the boundaries of the 4UY2 structurally determined ligand-binding domain of the receptor. The aligned nAchR putative orthologs a median of 83% amino acid percent identity (PID) among all sequence distances, 71% PID between the most distant two sequences in the set, a median of 94% PID among all species in set when considering only positions within 8Ang of contact with alpha-neurotoxin binding site, and when excluding fishes, 97% PID nearly completely conserved across mammals, avians, amphibians, and reptiles. To calculate conservation at each structural position, the sequences were rendered non-redundant at 99 % ID, leaving 85 unique sequences. A positional weight matrix of amino acid frequencies at each position was calculated and a Simson's diversity index diversity calculation was performed on each position.

Purification for Crystallography

Alpha-elapitotoxin-Dpp2a (3L24_DENPO) from Dendroaspis polylepis (black mamba), alpha-cobratoxin/long neurotoxin 1 (3L21_NAJNI) from Naja nivea (Cape cobra), Long neurotoxin 1 (3L21_OXYSC) from Oxyuranus scutellatus scutellatus (coastal taipan), and Alpha-delta bungarotoxin (3L2A_BUNCE) from Bungarus caeruleus (common krait) and antibody Centi-3FTX-D09 genes were synthesized and subcloned into pVRC8400 vectors, with HRV3C cleavable His or Fc tag (GenScript, NJ). Centi-3FTX-D09 IgG plasmids (Heavy and light chain) were transfected into Expi293F cells (Thermo Fisher) at 1:1 ratio (0.5mg heavy chain plasmid and 0.5mg of light chain plasmid per liter of cell culture), while long neurotoxins were transfected into 293 Freestyle cells (Thermo Fisher). Turbo293 transfection reagent (Speed BioSystems) was used according to manufacturer’s protocol at a ratio of 1mg of plasmid to 3ml of transfection reagent. Cells were cultured at 37 °C, 8% CO_2_ and 80% humidity for six days. Culture supernatants were harvested by centrifugation, and the resulting supernatants were filtered through 0.2um membranes. Proteins were affinity purified with protein A resin (GE) for Centi-3FTX-D09 IgG or cOmplete™ His-Tag Purification Resin (Roche) for long neurotoxin. His-tag and Fc region were cleaved by digestion with HRV3C protease (produced in house) followed by a second step of affinity purification with protein A resin to remove Fc fragment and uncleaved IgG or Ni-resin to remove cleaved his tag.

Fab fragment for Centi-3FTX-D09 antibody was further purified by size exclusion chromatography on a Superdex200 column (Cytiva) in PBS buffer. The peak corresponding to Fab fragment was concentrated to 10 mg/ml using a spin concentrator with MWCO of 10kDa and used to form complex with long neurotoxins.

Krait long neurotoxin (alpha-bungarotoxin) was also purchased from BioTechne (Catalog # 2133) in lyophilized form. Protein was dissolved in PBS and purified by size exclusion chromatography (Superdex 200, Cytiva). Peak corresponding to toxin was pooled and concentrated to 1mg/ml by spin concentration on 3kDa cutoff membranes.

Biolayer interferometry reactivity to whole venom

Affinity of long neurotoxins to Centi-3FTX-D09 -Fab was assessed using a fortéBio Octet instrument. His-tagged long neurotoxins were immobilized on Ni-NTA biosensors, then dipped into Centi-3FTX-D09 -Fab in a 2-fold concentration series. Sensograms of the concentration series were corrected with corresponding blank curves and fitted globally with Octet evaluation software using a 1:1 Langmuir model of binding.

Crystallization

Each neurotoxin was mixed with Centi-3FTX-D09 Fab fragment at 1.5:1 molar ratio (Toxin:Fab) and incubated at 4 °C overnight. Excess free toxin was removed by separation on Superdex 200 column (Cytiva). Peak corresponding to complex was pooled and concentrated to 5 mg/ml using spin concentrator (MWCO 30kDa). Complex between toxin and Fab was confirmed by SDS-PAGE and used immediately for crystallization trials.

Crystallization conditions were screened using Hampton Research, Wizard, and QIAGEN crystal screening kits. Crystal trays were set up using a Mosquito crystallization robot. Crystals initially observed from the wells were manually reproduced. Crystals suitable for data collection were obtained from the following conditions: Taipan-Centi-3FTX-D09: 100mM sodium acetate pH 5.5, 14% PEG8000, 2.65% PEG 400, 500mM NaCl, 10mM Mg2Cl; Mamba- Centi-3FTX-D09: 0.1M Tris pH 8.5, 10% PEG4000, 50mM Proline; Cobra-3FTX-D09: 200 mM (NH4)2SO4, 100 mM sodium acetate pH 4.6, 19% PEG 4000; Krait-Centi-3FTX-D09: 40% PEG 400, 5% PEG 3350, 0.1M sodium acetate pH 5.5.

Data collection and processing

For data collection the crystals were briefly dipped into reservoir solution supplied with 30% glycerol and flash-frozen in liquid nitrogen. X-ray diffraction data were collected at SER-CAT station 22-ID (Advanced Photon Source, Argonne National Laboratory, Argonne, IL). Prior to data collection the station was tuned up to the wavelength of 1A, the size of the beam was set to 50 um and the crystal was equilibrated to 100K with dry nitrogen stream. The data was collected using continuous rotation method. To achieve completeness and multiplicity 720 degrees of data were collected. Registered intensities were indexed, scaled, and merged with HKL2000 (*8*). Data collection statistics is shown in the **Table S4**.

Structure determination and analysis

The structures were solved by molecular replacement method with BALBES (*9*). Initial solution was verified and manually corrected with COOT (*10*). The coordinates were refined with REFMAC5 (*11*) (CCP4 suite (*12*)) and PHENIX.REFINE (*13*) (PHENIX suite (*14*)) alternating with manual revision of the model with COOT. Structure validation was performed with the Protein Data Bank validation server. Refinement statistics is provided in the **Table S4**. The coordinates and structure factors for the Taipan-Centi-3FTX-D09, Mamba- Centi-3FTX-D09, Cobra- Centi-3FTX-D09and Krait- Centi-3FTX-D09 complexes were deposited to the Protein Data Bank, and are available under accession codes 8D9Y, 8DA0, 8D9Z, and 8DA1, respectively.

Interface analysis for all toxin-fab complexes and for the acetylcholine receptor-Krait 3FTX (PDB ID: 4UY2 (*15*)) was performed with PISA (*16*). The details are presented in **Table S5 and S6**. The figures illustrated the structures and the Fab/nAChR-3FTX intermolecular interactions were generated by PYMOL (*17*).

Sequence variation was calculated for each residue as normalized Shannon's entropy based on 3FTX diversity analysis alignment (gaps were penalized with p(x_gap_)*log(1/n) and mapped on Krait structure – and colored purple to white to indicate diversity to conservation among sequences.

In vivo

All animal husbandry and experimental protocols were conducted under the guidance of approved IACUC animal use protocol CR-0119 reviewed, approved and assigned by Charles River D Laboratory (CRADL, South San Francisco, CA) IACUC administrator.

The studies used 8-week old C57BL/6 mice with body weights ranging between 18-22g. Mice were maintained in plastic boxes with water and food ad libitum, with 12 hour dark/light cycles.

Lyophilized venoms were obtained from multiple sources, rehydrated, aliquoted, frozen and thawed just prior to administration. Median lethal dose (LD_50_) and Maximal lethal dose (LD_100_) of each recombinant a-neurotoxin and whole venom were determined by an animal-sparing “up-and-down” methodology, with core body temperature as an adjunct to endpoint determination. The animal-sparing “up-and-down” methodology of estimating the LD_50_ that involved serial intradermal injection of predetermined mg/kg dosages of venom. Cates CC, McCabe JG, Lawson GW, *et al*.

For all in vivo challenges, venom and/or toxin was injected intraperitoneally and monitored via temperature and body weight every 20 minutes for 2-4 hours and every 8 hours sequentially, for 72 hours total. Treatments (Centi-3FTX-D09) was mixed with venom and/or toxin 30 minutes prior to injection. Varespladib (small molecule inhibitor of PLA2) was formulated in polyethylene glycol-300 (PEG) and Tween 80, and for its treatment group was injected 20 minutes prior to challenge injection. In accordance with the IACUC protocol, animals were monitored and mice will be euthanized if distress signs appear, including body temperature reduction greater than 4C, bodyweight loss more than 20% in 24 hours, and hunched posture, ruffled fur, prolonged lack of movement, moribundity, bleeding, or no signs of reversibility. Euthanization was by C02 followed by cervical dislocation.

In the in vivo protection assays, a challenge dose of 1xLD09_0-100_ of a venom or purified toxin was mixed with 30mg/kg of Centi-3FTX-D09 (diluted in PBS) and pre-incubated at 25°C for 30 minutes. The mixture containing 1x LD09_0-100_s of venom, were injected intraperitoneally into mice.

Statistical methods

Kaplan–Meier curves were plotted for both *Naja naja* venom LD_50_ and LD_100._ Log-rank test was used to examine the difference of survival curves between control and treatment groups. Survival rate was also compared based on a two-sample proportion test for venom LD_50._ Median survival time was calculated for both control and treatment groups in venom LD_50._ Analyses were performed using R version 3.3.3.

1.	K. M. Hastie, H. Li, D. Bedinger, S. L. Schendel, S. M. Dennison, K. Li, V. Rayaprolu, X. Yu, C. Mann, M. Zandonatti, R. Diaz Avalos, D. Zyla, T. Buck, S. Hui, K. Shaffer, C. Hariharan, J. Yin, E. Olmedillas, A. Enriquez, D. Parekh, M. Abraha, E. Feeney, G. Q. Horn, CoVIC-DB team1, Y. Aldon, H. Ali, S. Aracic, R. R. Cobb, R. S. Federman, J. M. Fernandez, J. Glanville, R. Green, G. Grigoryan, A. G. Lujan Hernandez, D. D. Ho, K.-Y. A. Huang, J. Ingraham, W. Jiang, P. Kellam, C. Kim, M. Kim, H. M. Kim, C. Kong, S. J. Krebs, F. Lan, G. Lang, S. Lee, C. L. Leung, J. Liu, Y. Lu, A. MacCamy, A. T. McGuire, A. L. Palser, T. H. Rabbitts, Z. Rikhtegaran Tehrani, M. M. Sajadi, R. W. Sanders, A. K. Sato, L. Schweizer, J. Seo, B. Shen, J. L. Snitselaar, L. Stamatatos, Y. Tan, M. T. Tomic, M. J. van Gils, S. Youssef, J. Yu, T. Z. Yuan, Q. Zhang, B. Peters, G. D. Tomaras, T. Germann, E. O. Saphire, Defining variant-resistant epitopes targeted by SARS-CoV-2 antibodies: A global consortium study. *Science*. **374**, 472–478 (2021).

2.	E. Rujas, I. Kucharska, Y. Z. Tan, S. Benlekbir, H. Cui, T. Zhao, G. A. Wasney, P. Budylowski, F. Guvenc, J. C. Newton, T. Sicard, A. Semesi, K. Muthuraman, A. Nouanesengsy, C. B. Aschner, K. Prieto, S. A. Bueler, S. Youssef, S. Liao-Chan, J. Glanville, N. Christie-Holmes, S. Mubareka, S. D. Gray-Owen, J. L. Rubinstein, B. Treanor, J.-P. Julien, Multivalency transforms SARS-CoV-2 antibodies into ultrapotent neutralizers. *Nat. Commun.* **12**, 3661 (2021).

3.	UniProt Consortium, UniProt: the universal protein knowledgebase in 2021. *Nucleic Acids Res.* **49**, D480–D489 (2021).

4.	J. Glanville, T. C. Kuo, H.-C. von Büdingen, L. Guey, J. Berka, P. D. Sundar, G. Huerta, G. R. Mehta, J. R. Oksenberg, S. L. Hauser, D. R. Cox, A. Rajpal, J. Pons, Naive antibody gene-segment frequencies are heritable and unaltered by chronic lymphocyte ablation. *Proceedings of the National Academy of Sciences*. **108**, 20066–20071 (2011).

5.	E. W. Sayers, E. E. Bolton, J. R. Brister, K. Canese, J. Chan, D. C. Comeau, R. Connor, K. Funk, C. Kelly, S. Kim, T. Madej, A. Marchler-Bauer, C. Lanczycki, S. Lathrop, Z. Lu, F. Thibaud-Nissen, T. Murphy, L. Phan, Y. Skripchenko, T. Tse, J. Wang, R. Williams, B. W. Trawick, K. D. Pruitt, S. T. Sherry, Database resources of the national center for biotechnology information. *Nucleic Acids Res.* **50**, D20–D26 (2022).

6.	HMMER. *hmmer.org*, (available at www.hmmer.org).

17.	PyMOL, (available at http://www.pymol.org/pymol).

18.	J. Glanville, W. Zhai, J. Berka, D. Telman, G. Huerta, G. R. Mehta, I. Ni, L. Mei, P. D. Sundar, G. M. R. Day, D. Cox, A. Rajpal, J. Pons, Precise determination of the diversity of a combinatorial antibody library gives insight into the human immunoglobulin repertoire. *Proceedings of the National Academy of Sciences*. **106**, 20216–20221 (2009).
