## Supplementary material for "Venom protection by broadly neutralizing antibody from a snakebite subject": Ethics statement

Ethics Statement for

**This PDF file includes:**

Ethics Statement

Ethics Statement

The study of human samples was strictly non-interventional: the subject was not directed to expose himself to venom nor was the subject provided any guidance on venom self-exposure. Over a 17-year period prior to interacting with researchers, subject had independently chosen to self-immunize with snake venom, and during this time the subject had independently developed a standard protocol for periodic venom self-immunizations. Subject notified researchers of subject’s self-immunization schedule, and researchers arranged for a commercial phlebotomy service to obtain the blood samples before and 14 days following the subject’s next self-scheduled self-immunization. Informed consent was obtained for the two 20ml blood samples collected. Collection was conducted in accordance with Western Institutional Review Board IRB Exemption Work Order #1-1209200-1. No further samples were taken. Researchers provided materials to subject on the risks of the self-envenoming. In 2018, subject retired from self-envenoming.

All animal husbandry and experimental protocols were conducted under the guidance of approved IACUC animal use protocol CR-0119, reviewed, approved and assigned by Charles River D Laboratory (CRADL, South San Francisco, CA) IACUC administrator. To minimize number of animals used, an animal-sparing “up-and-down” methodology of estimating the LD_50_ that involved serial intradermal injection of predetermined mg/kg dosages of venom. Cates CC, McCabe JG, Lawson GW, *et al*. In accordance with the IACUC protocol, animals were monitored and mice will be euthanized if distress signs appear, including body temperature reduction greater than 4C, bodyweight loss more than 20% in 24 hours, and hunched posture, ruffled fur, prolonged lack of movement, moribundity, bleeding, or no signs of reversibility. Euthanization was by C02 followed by cervical dislocation.
